# Brain-wide mapping and synaptic localization of C1QL3 using a novel epitope-tagged knock-in mouse

**DOI:** 10.64898/2026.03.05.709958

**Authors:** William P. Armstrong, James M. Salvatore, Matthew J. Sticco, Keaven Caro, J. Wesley Maddox, Angie Huang, Brenna C. McAllister, Christopher O’Connell, Siu-Pok Yee, Amy Lee, Susanne Ressl, David C. Martinelli, Alexander C. Jackson

## Abstract

Synapse formation and function are coordinated spatially and temporally by a host of synaptic proteins that regulate neuronal signaling, synapse specificity, and plasticity; many of which are implicated in neuropsychiatric disorders. Members of the C1q/TNF superfamily function as synaptic organizers, shaping synapse assembly and maintenance. Among them, C1QL3 plays a putative role in trans-synaptic adhesion and modulation of synaptic strength, but the lack of a reliable antibody to detect it has severely limited the ability to map its endogenous localization and study its biochemical properties. Here, we present a novel epitope-tagged knock-in mouse line (*C1ql3*^2HA^), in which two hemagglutinin (HA) epitopes were inserted near the N-terminus of the endogenous C1QL3 protein. This model enables purification, detection, and subcellular localization of native C1QL3 protein (C1QL3-2HA) with high specificity, eliminating the need for overexpression or custom antibodies. We validated that *C1ql3*^2HA^ mice maintain normal mRNA expression, biochemical properties, and behavior. Using native PAGE, we determined the endogenous oligomeric state of C1QL3-2HA. Brain-wide light-sheet microscopy uncovered an expanded neuroanatomical map of C1QL3-2HA expression, including newly identified populations in cortical and subcortical regions as well as the retina. Dual immunohistochemistry confirmed cell type-specific expression patterns, and super-resolution STED microscopy localized C1QL3-2HA to hippocampal mossy fiber synapses, positioned between pre- and post-synaptic markers, supporting its hypothesized role in trans-synaptic complexes. This knock-in mouse line is a powerful tool for studying the anatomical, molecular, and synaptic biology of C1QL3 in all cellular/tissue contexts, enabling future studies into its potential roles in the nervous system and beyond.

## INTRODUCTION

Neuronal circuits are built and remodeled through the coordinated actions of synaptic organizers - secreted or membrane-anchored proteins that guide synapse formation, target specificity, synaptic signaling, and plasticity (Connor & Siddiqui, 2023; Südhof, 2021; Yuzaki, 2018). Members of the complement component 1, q subcomponent/tumor necrosis factor (C1q/TNF) superfamily, including the complement component 1, q subcomponent-like (C1QL) proteins, have emerged as critical regulators of synaptic structure and function across the central nervous system (CNS) and beyond (Carland & Gerwick, 2010; Ghai et al., 2007; Matsuda, 2017; Peña Palomino et al., 2023; Südhof, 2023; Yuzaki, 2010, 2018). These proteins mediate trans-synaptic adhesion, signaling, and the stabilization of synaptic contacts.

A member of this family, C1QL3, has garnered increasing attention owing to its strong CNS enrichment and high degree of evolutionary conservation. C1QL3 (also known as CTRP13 and identified initially as K100) was first cloned in a yeast two-hybrid screen using heat shock protein 47 (HSP47), a collagen-specific ER chaperone, as bait (Koide et al., 2000). Its C-terminal globular C1q domain (gC1q) is 100% identical between human and mouse and alone is sufficient to mediate all currently known C1QL3-protein interactions. C1QL3 interacts with key synaptic binding partners, including the adhesion G protein-coupled receptor B3 (ADGRB3, or BAI3), specific kainate receptor paralogs, specific neurexin-3 splice variants, and neuronal pentraxins, placing it at the center of multiple trans-synaptic pathways involved in synapse regulation (Bolliger et al., 2011; Matsuda et al., 2016; Miao et al., 2025; Sticco et al., 2021). Despite this, the precise endogenous protein distribution in the brain, subcellular localization, and molecular assembly state of C1QL3 remain largely unexplored.

Although C1QL3 is also expressed in select peripheral tissues (*e.g.,* adipose tissue) and has been implicated in regulating systemic metabolism, insulin secretion, and lipid/glucose homeostasis (Byerly et al., 2013; Chen et al., 2023; Gupta et al., 2018; Koltes et al., 2019; Wei et al., 2011), its most prominent expression is in the CNS (Iijima et al., 2010). Within the brain, *in situ* hybridization and mRNA reporter analyses have demonstrated expression in discrete neuronal populations, including pyramidal cells of the cerebral cortex, hippocampus, and amygdala, as well as select subcortical brain regions such as the prominent expression in the suprachiasmatic nuclei (Chew et al., 2017; Iijima et al., 2010; Martinelli et al., 2016). Work on its paralogs, C1QL1 and C1QL2, has further revealed that proteins of this family localize to excitatory synapses, where they contribute to synapse formation, adhesion, synaptic signaling, and regulation of behavior (Aimi et al., 2023; Biswas et al., 2021; Caro et al., 2025; Chew et al., 2017; Kakegawa et al., 2015, 2024; Koumoundourou et al., 2024; Martinelli et al., 2016; Matsuda et al., 2016; Pan et al., 2024; Paul et al., 2025; Sigoillot et al., 2015; Wang et al., 2020; Zhang et al., 2025).

We and others previously characterized *C1ql3* gene expression using a mVenus reporter allele (*C1ql3*^flox-mVenus^) and *in situ* hybridization (Chew et al., 2017; Iijima et al., 2010; Martinelli et al., 2016), but both approaches have limitations: they report mRNA rather than protein and do not resolve subcellular protein localization. Progress on studying C1QL proteins has been slowed by a dearth of commercially available antibodies for immunolocalization and biochemistry, which typically only work to detect overexpressed proteins. To address these challenges, we created a *C1ql3* knock-in (KI) mouse allele in which a double hemagglutinin (2HA) epitope tag was inserted immediately downstream of the endogenous signal peptide. This strategy enables detection of native C1QL3 protein expression, using widely available anti-HA antibodies.

We verified that the *C1ql3*^2HA^ allele was correctly targeted and did not disrupt *C1ql3* mRNA expression or gross behavioral phenotypes. We further demonstrated that the HA-tagged C1QL3 protein (C1QL3-2HA) can be purified from mouse brain tissue, and for the first time, showed that endogenous C1QL3 forms defined higher-order oligomers, consistent with biochemical findings from recombinant systems (Sticco et al., 2021). Using whole-brain light-sheet microscopy, we created a comprehensive neuroanatomical atlas of C1QL3 expression across the brain and retina, revealing many previously undetected populations, including ones in the cerebral cortex, thalamus, midbrain, hypothalamus, brainstem, and cerebellum. Dual immunostaining confirmed the expression of C1QL3 in specific types of excitatory and inhibitory neurons. Finally, using super-resolution stimulated emission depletion (STED) microscopy, we localized C1QL3-2HA protein within the putative synaptic cleft of hippocampal mossy fiber synapses, between pre- and post-synaptic markers, consistent with its hypothesized role as a secreted trans-synaptic organizer (Martinelli et al., 2016; Matsuda et al., 2016).

Together, our findings provide a comprehensive anatomical, molecular, and subcellular profile of C1QL3 in the mouse CNS. This *C1ql3*^2HA^ KI mouse line establishes a valuable new tool for studying synaptic organization and C1QL3-mediated adhesion mechanisms in both normal brain function and disease. It also joins a growing family of successful epitope-tagged mouse models for mapping synaptic proteins that lack sufficient commercially available antibody options (Cheung et al., 2024; Lloyd et al., 2023; Nozawa et al., 2018; Trotter et al., 2019; Vyas et al., 2020).

## METHODS

This study was carried out in strict accordance with the recommendations in the Guide for the Care and Use of Laboratory Animals of the National Institutes of Health. The protocols were approved by the University of Connecticut Institutional Animal Care and Use Committee (IACUC; protocols 201026-0926 and A22-056). Mice were housed in the standard 12-hour light, 12-hour dark cycle and fed standard chow. All procedures were performed under ketamine/xylazine anesthesia, and all efforts were made to minimize suffering.

### Creation of *HA-C1ql3* allele

Mice were generated by electroporation of one-cell embryos with ESP Cas9 protein (150 ng/µl; Sigma, cat# ESPCAS9PRO); sgRNA (50 ng/µl; IDT) and ssODN (100 ng/µl; IDT). Sequence of guide RNA: 5’- CTCGTAGTGAGCCGACGTGC (antisense).

Sequence of ssDNA donor DNA: 5’- G*T*G* ATG GTG CTT CTG CTG GTC ATC CTC ATC CCG GTG CTG GTG AGC TCG GCT GGC ACG TCG GCT GCT AGC TAC CCC TAC GAT GTG CCC GAT TAC GCC GGA GGA TAC CCC TAC GAT GTG CCC GAT TAC GCC GGC GCT AGC CAC TAC GAG ATG CTG GGC ACC TGC CGC ATG GTC TGC GAC CCC TAT GGG GGC ACC AAG GC*T* C*C -3’, * represents phosphorothioate and is designed to minimize any degradation due to exonuclease. Following electroporation, embryos were transferred to pseudopregnant females. For genotyping primers: *HA-C1QL3*-F 5’-TGATCGCCGCCGGGGCGCTG and *HA-C1QL3*-R 5’- GTCCGGAGTGGCGGCGGTGC. The PCR program protocol was 94°C for 3 min, 94°C for 15 sec, 68°C for 15 sec, 72°C for 15 sec, and steps 2 – 4 were repeated 34 times. The knock-in band was 248 bp whereas the WT band was 173 bp long. The polymerase used was Denville Choice Taq Mastermix (cat. # 1001111). We sequenced the entire open reading frame of exon 1 and no mutations were observed in this amplicon. The line was backcrossed to WT C57BL/6J mice for 4 generations before creating homozygotes.

### Denaturing PAGE

Recombinant C1QL3-2HA obtained by transfecting HEK-293T cells with pDisplay C1QL3-2HA plasmid (Bolliger et al., 2011) or pCMV5 2xHA-C1ql3 (cloned for this manuscript) and harvesting from conditioned media. Cerebral cortex tissue and blood samples were harvested from adult *C1ql3^2HA^* mouse and a wild type litter mate. Blood samples were collected in syringes coated with 1 unit/µl heparin and stored in 1.5 ml tube with heparin to final concentration of 20 units of heparin/ml of blood. After centrifugation, plasma supernatant collected then denatured for Western blot. Brain tissue pieces were homogenized with a Dounce homogenizer in RIPA buffer plus protease inhibitors. The samples were centrifuged for 40 min at 14.8K RPM and supernatants were then denatured for Western blot using a PVDF membrane. Blot probed with 1:500 rat anti-HA (clone 3F10, Roche, Switzerland). The secondary antibody, 1:3000 goat anti-rat, HRP (R&D Systems, #HAF005) applied followed by standard ECL detection (refer to the antibody list in Table 1).

**Table 1:**
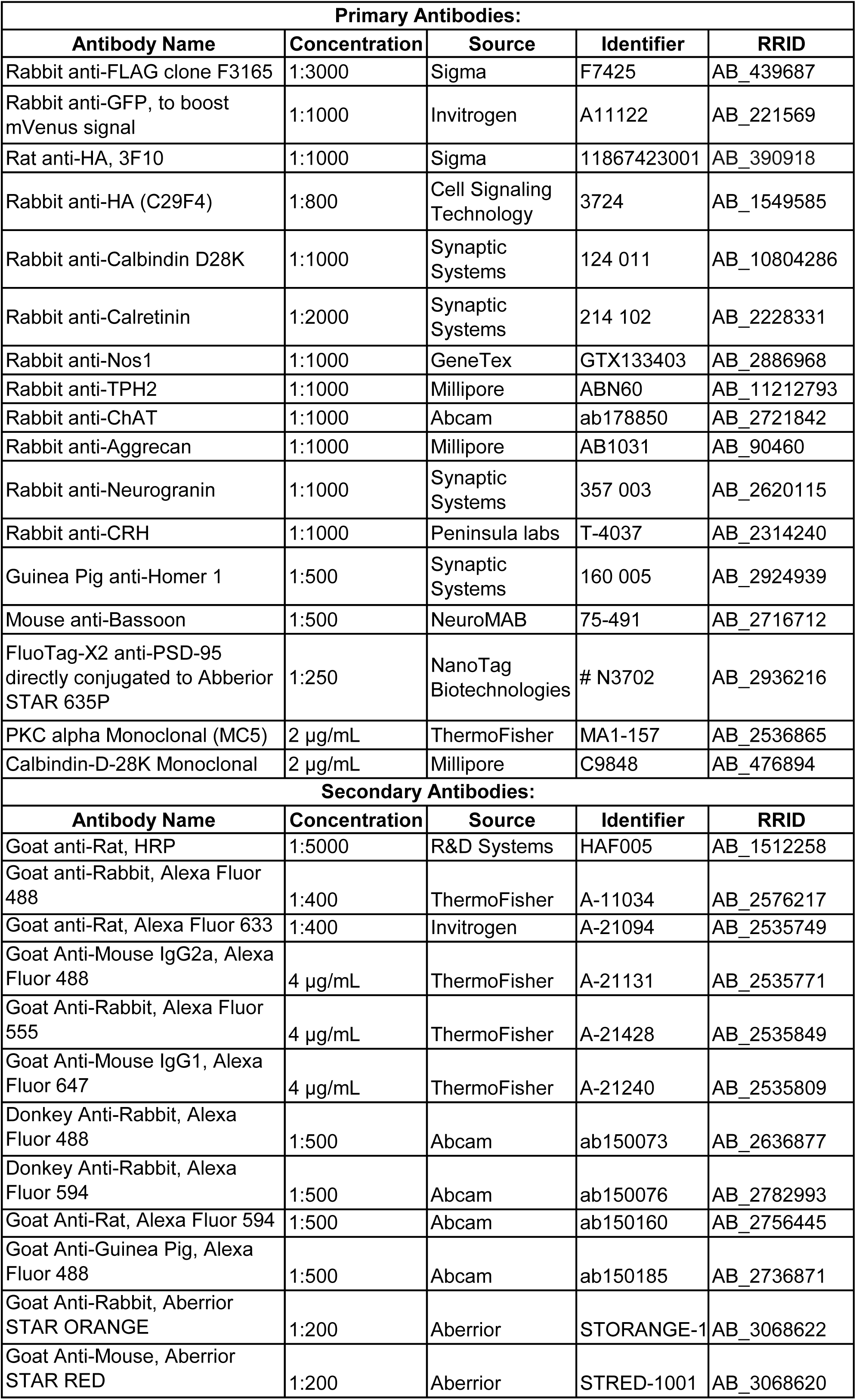
List of primary and secondary antibodies used.

### Blue Native PAGE

A *C1ql3^2HA^* brain was homogenized with a Dounce homogenizer in 10 ml of solubilization buffer (1% Triton X-100, 150 mM NaCl, 25 mM HEPES, 2 mM CaCl_2_. pH 7.4, protease inhibitors). Lysate was rocked for 30 min at 4°C then centrifuged at 4500 RPM for 30 min at 4°C. The cleared lysate was incubated overnight while mixing at 4°C with 100 µl monoclonal anti-HA−agarose (A2095 Sigma, USA). The affinity matrix containing the bound proteins was transferred to Screw Cap Spin Columns (ThermoScientific Pierce, USA) and washed a total of 5 times with buffer (same as solubilization buffer but no detergent/inhibitors). Acid elution was performed twice with 0.5 ml 100 mM glycine, 150 mM NaCl, pH 2.5, and then pH was normalized with 60 µl 3 M TRIS, pH 8.0. Buffer exchange to the wash buffer performed using Amicon Ultra 0.5 ml Centrifugal Filter Unit, 30K (Cat# UFC503024). For the PAGE, ∼15 µl of purified protein was mixed with Invitrogen NativePAGE 4x Sample Buffer (BN2003, Invitrogen, USA) and loaded onto a NativePAGE 3 to 12%, Bis-Tris gel (BN1001BOX, Invitrogen, USA) using a Mini Gel Tank (A25977, Invitrogen, USA). Run side-by-side was 1 ng of C1QL3-2HA produced from HEK293T cells. For molecular weight estimation, we used NativeMark Unstained Protein Standard (LC0725, Invitrogen, USA). Electrophoresis was performed according to the instructions for the NativePAGE gel, using standard anode buffer and light blue cathode buffer containing Coomassie G-250. The protein standard was visualized with routine staining with Coomassie Blue R-250, while rest of gel was transferred to PVDF membrane with a Trans-Blot Turbo (Bio-Rad, USA) and NuPAGE Transfer Buffer (NP0006, Invitrogen, USA), followed by Western blot as described above.

### Cell surface binding assay

Recombinant protein production/purification and *in vitro* cell surface binding assay performed exactly as described previously (Sticco et al., 2021). In brief, the previously used expression vectors were transiently transfected into COS-7 cells with the calcium phosphate method onto glass coverslips. 2 days later, cells washed with room temperature binding buffer containing NaN_3_ (azide blocks endocytosis). Purified exogenous C1QL3-2HA applied to cells at 150 nM for 60 min at room temp. Cells were washed 4 times then fixed for 10 min with 4% PFA. Cells blocked with 5% goat serum in PBS for 1 hour, followed by indirect immunofluorescence using anti-HA and anti-flag antibodies and appropriate secondary antibodies. To quantify, 15 healthy appearing transfected cells (determined by large and flat morphology) were selected and photographed at random. The magnetic lasso tool was used in Adobe Photoshop CSS extended version 12.0 to outline the cell’s plasma membrane, then the mean fluorescence intensity value acquired.

### Behavioral analysis

Spontaneous locomotor activity was measured using a PAS Open Field system (San Diego Instruments), as described in (Ma et al., 2008). Animals were placed in the center of a 15” × 15” chamber for 15 minutes and their ambulatory activity was recorded by photocell beam breaks.

### qRT-PCR

Total RNA was harvested from freshly harvested cerebral cortices from 8-week old *C1ql3^2HA^* and wild type littermate mice using TRIzol phenol:chloroform extraction. RNA was reverse transcribed using Denville rAmp cDNA synthesis kit (C188R97). Primers used were: *C1ql3* Forward CTACTTCTTCACCTACCACGTC, *C1ql3* Reverse GAAGGACCACACTGTTACTGG, *Gapdh* Forward ATGACATCAAGAAGGTGGTG, *Gapdh* Reverse CATACCAGGAAATGAGCTTG, identical to those used previously (Martinelli et al., 2016). Each cDNA sample (100 ng) was added to a master mix containing primers for either *C1ql3* or *Gapdh* and BioRad SsoAdvanced universal SYBR green supermix (cat #1725270). Reactions were performed on a BioRad CFX96 Real-Time PCR machine (BioRad). Quantification was relative to *Gapdh*.

### Brain immunofluorescence

Adult mice (2-4 months old) were anesthetized with ketamine/xylazine intraperitoneally and transcardially perfused with saline followed by 4% paraformaldehyde (PFA) in PBS. For typical immunohistochemistry, dissected brains were post-fixed in 4% PFA for 24 hours, cryoprotected in 30% sucrose for 48 hours, and flash-frozen in freezing isopentane. Coronal brain cryosections (40 μm) were washed in PBS and PBST (0.2% tween in PBS) before blocking for 2 hours in 2% normal donkey serum. Sections were then incubated in primary antibodies overnight, then washed with subsequent PBST before 2 hours of incubation in secondary antibodies (refer to antibody list in Table 1). Sections were mounted onto slides (Cat. # 125442, Fisher Scientific, USA) with coverslips (Ref. # 12460S, Epredia, Germany) using Prolong Gold mounting media with DAPI (Ref. # P36931, Invitrogen, USA).

Fluorescence images were taken on either a Keyence fluorescence microscope (BZ-X700) or a Leica SP8 laser confocal microscope through the University of Connecticut Advanced Light Microscopy Facility, (RRID:SCR_027547).

### Tissue preservation and clearing, immunolabeling and imaging (light-sheet)

Paraformaldehyde-fixed tissue samples were preserved with SHIELD reagents (Park et al., 2019) according to the manufacturer’s protocol (LifeCanvas Technologies, Cambridge, MA). Samples were subsequently delipidated using Clear+ delipidation reagents (LifeCanvas Technologies). Following delipidation, tissues were blocked in Antibody Blocking Solution (LifeCanvas Technologies) with 5% normal donkey serum (Jackson ImmunoResearch Laboratories Inc). Immunolabeling was performed using the SmartBatch+ device and the RADIANT Buffer system (LifeCanvas Technologies), incorporating stochastic electrotransport (Kim et al., 2015) and eFLASH/CuRVE (Yun et al., 2025) technologies. Following immunolabeling, samples were incubated in 50% EasyIndex (diluted in DIW) overnight at 37°C, then transferred to 100% EasyIndex (RI = 1.52; LifeCanvas Technologies) for 24 hours for refractive index matching. After index matching, the samples were imaged using a SmartSPIM axially-swept light sheet microscope with a 3.6x objective (0.2 NA; LifeCanvas Technologies).

### Brain atlas registration

Samples were registered to the Allen Brain Atlas (Allen Institute: https://portal.brain-map.org/) using an automated process (alignment performed by LifeCanvas Technologies). A nuclear dye channel using oxazole yellow staining (Biotium, Cat# 152068-09-2) for each brain was registered to an average nuclear dye atlas (generated by LCT using previously registered samples). Registration was performed using successive rigid, affine, and b-spline warping algorithms (SimpleElastix: https://simpleelastix.github.io/).

### Cell detection

Automated cell detection was performed by LifeCanvas Technologies using a custom convolutional neural network created with the Tensorflow Python package (Google). The cell detection was performed by two networks in sequence. First, a fully-convolutional detection network (https://arxiv.org/abs/1605.06211v1) (Shelhamer et al., 2016) based on a U-Net architecture (https://arxiv.org/abs/1505.04597v1) (Ronneberger et al., 2015) was used to find possible positive locations. Second, a convolutional network using a ResNet architecture (https://arxiv.org/abs/1512.03385v1) (He et al., 2015) was used to classify each location as positive or negative. Using the previously calculated Atlas Registration, each cell location was projected onto the Allen Brain Atlas in order to count the number of cells for each atlas-defined region.

### Tissue preparation and immunostaining for STED microscopy

For STED microscopy, following cryoprotection, brains were embedded in OCT in cryomolds over freezing isopentane. Coronal brain cryosections (5 μm) were directly mounted onto #1.5 gelatin coated glass coverslips (Cat. # 72293-06, Electron Microscopy Sciences, USA). Sections were washed in distilled H_2_O for 5 minutes at 37°C and then in 1mg/ml pepsin in 0.2 N HCl at 37°C for antigen retrieval prior to PBS washes as states for typical IHC experiments. Coverslips were directly mounted onto glass slides with Prolong Gold mounting media without DAPI (Ref. # P36930, Invitrogen, USA). STED images were taken using an Abberior Instruments Expert Line STED microscope equipped with a 595 nm (for Alexa Fluor 488) and 775 nm (for STAR ORANGE and STAR RED) depletion lasers and an Olympus 100x/1.40 UPLANSAPO oil immersion objective, through the University of Connecticut Advanced Light Microscopy Facility, (RRID:SCR_027547). All analysis of STED imaging was performed using single planes to eliminate photobleaching following multiple z-steps confounding analysis.

### Image analysis

All line scans were conducted using the built-in FIJI ImageJ plugin “Line Profile” to measure fluorescence intensity through the cortex of light-sheet microscopy images or individual channels at CA3 synapses for STED images. For light-sheet images, a consistent 5 linear ROIs were drawn in the exact same region of each cortical subdivision using boundaries from image registration to the Allen Brain Atlas to denote layers 1-6. Fluorescence intensity across the line scan was normalized to average intensity for each scan to account for variable background between mice, and then the relative intensity through the cortex was averaged across mice. Prior to analysis, all STED images were deconvolved using Hyugens deconvolution software to eliminate background noise. Center-center distances between all BSN and HOMER-immunoreactive (IR) puncta in the 100X field of the STED images were calculated using Imaris (Oxford instruments). HA-IR puncta within the median BSN-HOMER distance from both BSN and HOMER were then analyzed to calculate the center-to-center distance between putative synaptic HA-IR puncta and either BSN or HOMER. For line scans of individual synapses, putative synapses with clear BSN-HOMER appositions that colocalized with HA-IR were selected. The same line scan extended through the entirety of each of the three puncta analyzed, and the same linear ROI was used for all three channels of the same synapse. Fluorescence intensity of each puncta type was plotted relative to distance from the peak of the HA-IR fluorescent signal and averaged in 0.025 µm bins across multiple putative synapses. Fluorescence intensity was normalized to peak intensity of each respective channel.

### Retina immunofluorescence

Eyes from adult mice 2-3 months old were fixed with 4% paraformaldehyde in 0.1 M phosphate buffer (PB), hemisected, lenses were removed, and eye cups were washed three times with PB containing 1% glycine followed by infusion of 30% sucrose in 0.1 M PB at 4°C overnight. The eye cups were orientated along their dorsal-ventral axis and frozen in a 1:1 (wt/vol) mixture of OCT compound and 30% sucrose in a dry ice/isopentane bath. Eye cups were cryosectioned at 20 µm on a Leica CM1850 cryostat (Leica Microsystems), mounted on Superfrost plus Micro Slides (VWR), dried for 5 to 10 min at 42°C, and stored at -20°C until used. All remaining steps were carried out at room temperature. Slides with mounted cryosections were washed with 0.1 M PB for 30 min to remove the OCT/sucrose mixture and blocked with dilution solution (DS, 0.1 M PB/10% goat serum/0.5% Triton-X100) for at least 30 min. Sections were incubated with primary antibodies (please refer the antibody list in Table 1) overnight and then washed five times with 0.1 M PB. Sections were then incubated with secondary antibodies (Table 1) for 30 min and then washed five times with 0.1 M PB. Trace 0.1 M PB was removed, and sections were then mounted with #1.5H coverslips (ThorLabs) using ProLong Glass Antifade Mountant with NucBlue (Thermo Fisher Scientific). Immunofluorescence in labeled retinal sections was visualized using an Olympus FV3000 confocal microscope (Tokyo, Japan) equipped with an UPlanApo 60x oil HR objective (1.5 NA). Images (1024 x 1024 pixels) were captured using the Olympus FLUOVIEW software package. Acquisition settings were optimized using a saturation mask to prevent signal saturation prior to collecting 16-bit. All confocal images presented are maximum z-projections.

## RESULTS

### Design and validation of a double HA epitope-tagged C1QL3 KI mouse

Our objective in generating a double HA epitope-tagged C1QL3 KI mouse was to effectively and reproducibly detect endogenously-expressed C1QL3 protein in the mouse brain. Single HA epitope-tagged recombinant C1QL3 (Figure 1A) showed no disrupted trafficking or binding and rescued a behavior phenotype caused by WT *C1ql3* KO (Martinelli et al., 2016; Sticco et al., 2021), and the availability of many excellent and validated anti-HA antibodies with various conjugation technologies made an HA tag for the KI C1QL3 mouse a promising strategy. Additionally, to increase sensitivity, we decided to create a double HA (2HA) tagged KI mouse.

**Figure 1:**
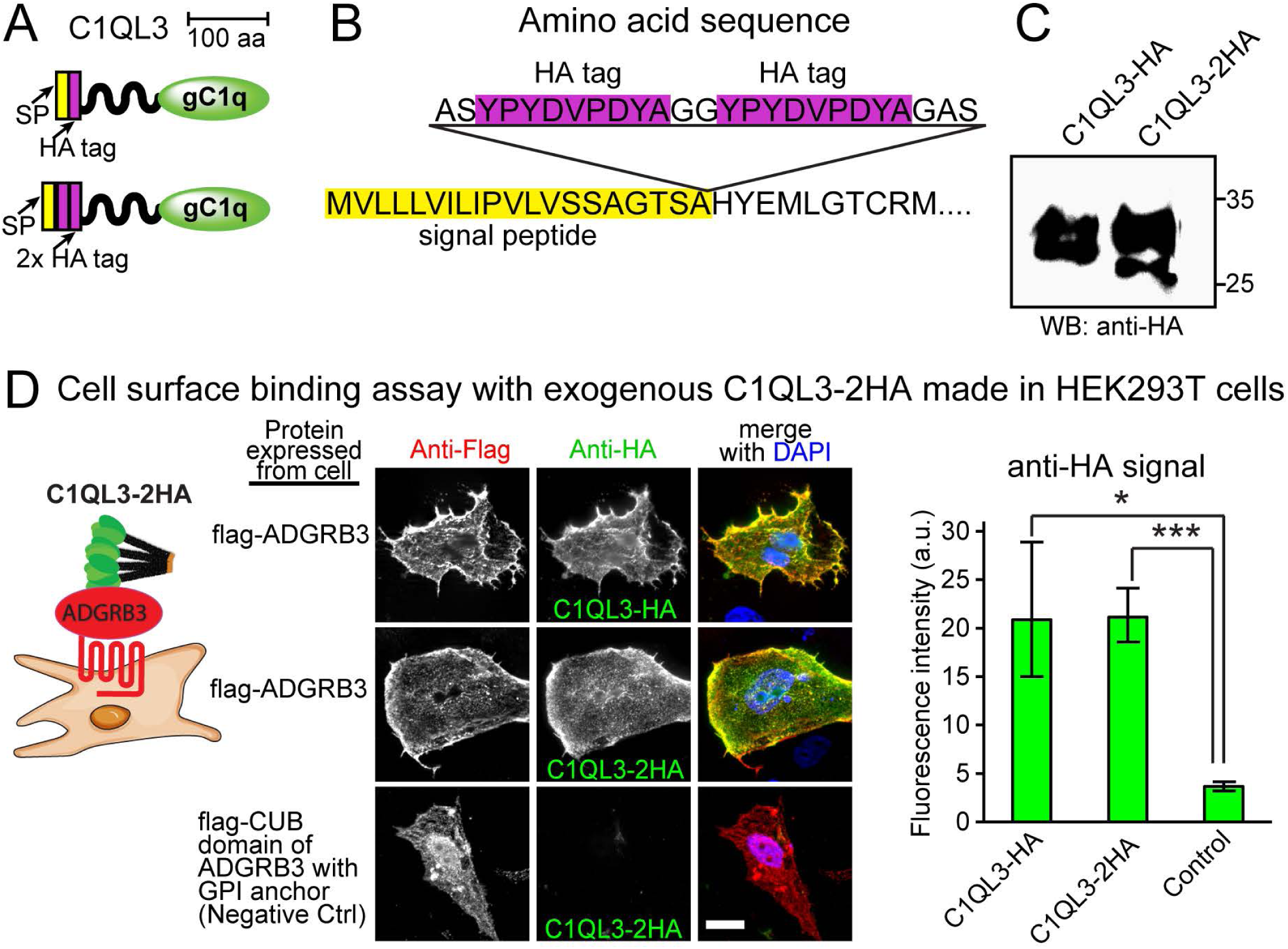
Validation of 2HA-tag design. **A)** Diagram of the location of the HA tags (violet) in relation to the signal peptide (SP, yellow) and gC1q domain (green). **B)** The amino acid sequence of the 2HA insertion (violet) following the SP (yellow). **C)** Denaturing PAGE and Western blot on the conditioned media from transiently transfected HEK-239T cells for each version of C1QL3. **D)** Cell surface binding assay performed on live cells with sodium azide to block endocytosis and equimolar amounts of 150 nM purified exogenous C1QL3. COS-7 cells were transfected to express either flag-tagged ADGRB3 or a GPI-anchored CUB domain of ADGRB3 that does not bind C1QL3 (Sticco et al., 2021). After fixation, immunofluorescence was performed without permeabilization to detect bound proteins. Immunofluorescence signal intensity of 15 randomly selected transfected cells is shown on the right. Statistical comparisons were made using one-way ANOVA with Bonferroni’s post hoc test (*P ≤ 0.05, ***P ≤ 0.001, scale bar=20 µm).

Therefore, before generating the mouse, we generated 2HA epitope-tagged recombinant C1QL3 and compared its binding affinity to the single HA-tagged C1QL3 (Figure 1A, B). When expressed from transiently transfected HEK-293T cells, each was found to be secreted equally into the media, albeit with some apparent differences in post-translational modifications (Figure 1C). In a live cell surface-binding assay, we assessed each purified C1QL3 for its retention on the surface of COS7 cells expressing ADGRB3. No differences were observed in the fluorescence intensity between C1QL3-HA- and C1QL3-2HA-immunoreactive (IR) signals on transfected cells (Figure 1D), suggesting that an additional HA epitope tag on C1QL3 is functionally inconsequential.

We then used CRISPR/Cas9 technology and the electroporation of one-cell embryos to target the double HA tag to the endogenous *C1ql3* locus. Targeting in the new KI mouse line (referred to as *C1ql3*^2HA^ hereafter) was confirmed by Sanger sequencing of genomic DNA (Figure 2A). Breeding of heterozygous parents produced homozygous KI pups in the approximate Mendelian ratio (Figure 2B), and *C1ql3*^2HA^ mice were viable and fertile. We detected HA-IR from brain tissue at the correct band size after denaturing PAGE and Western blot (Figure 2C). A slight difference in apparent size was observed when comparing this endogenously produced C1QL3-2HA to that produced in transfected HEK-293T cells, likely due to variations in post-translational glycosylation. Despite numerous reports of detectable C1QL3 in circulating blood (An et al., 2019; Bai et al., 2017; Fadaei et al., 2016; Shanaki et al., 2016; C. Wang et al., 2019; Wei et al., 2011), we did not observe any in plasma, although it is possible that it could be revealed with different methods.

**Figure 2:**
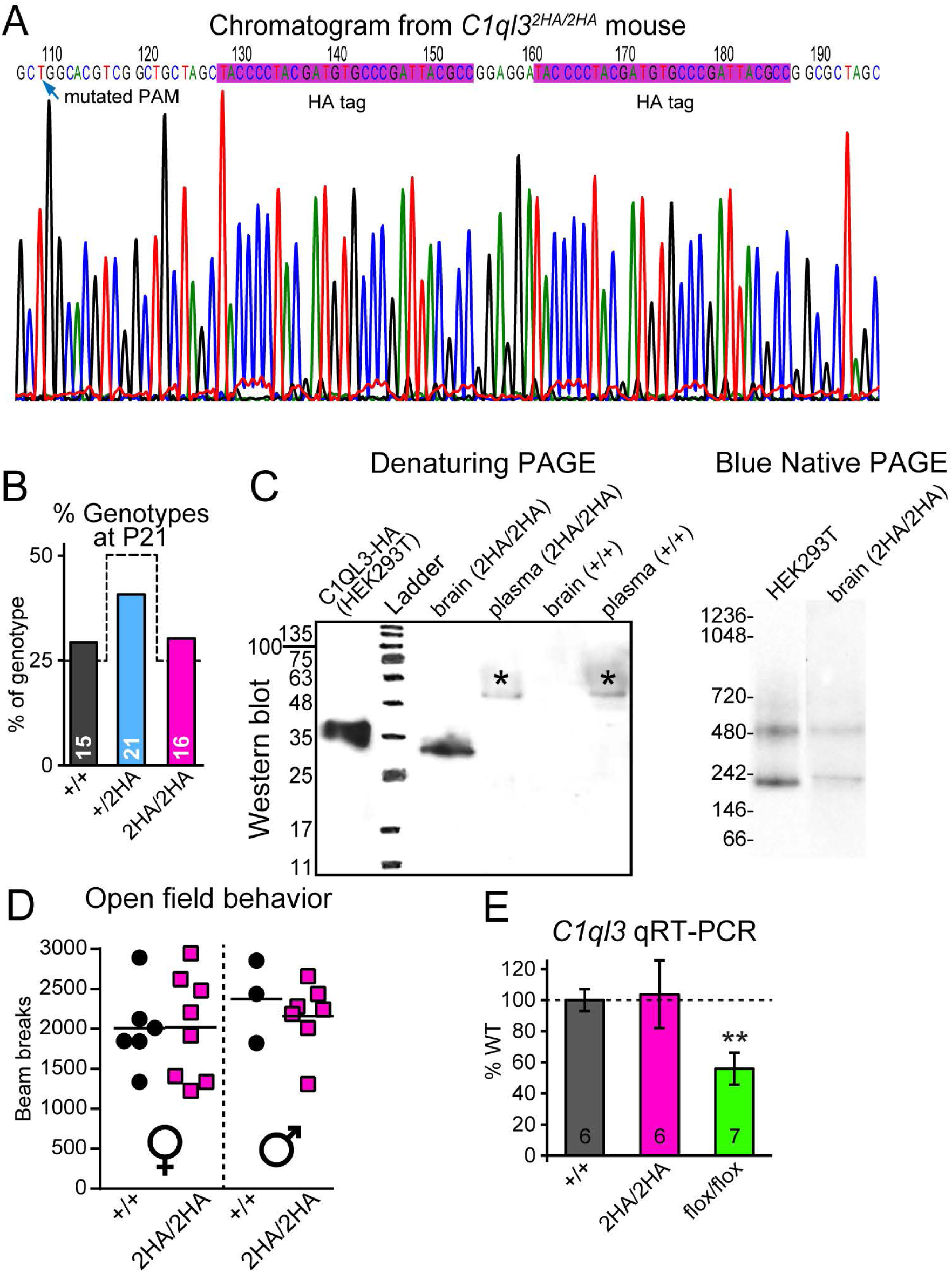
Generation of the novel 2HA-tagged allele. **A)** Chromatogram of the Sanger sequencing verification of the genomic DNA of a homozygous KI mouse. Note (blue arrow) that a synonymous/silent mutation was made to the PAM site to block further Cas9 targeting and prevent unwanted indels. **B)** The observed ratio of genotypes in offspring (7 litters) from heterozygous parents compared to the expected Mendelian ratio (dashed line). **C)** Left: Denaturing PAGE and anti-HA Western blot on dissected cerebral cortex homogenate or plasma (from blood) from homozygous KI (HA/HA) and WT (+/+) littermates, and positive control recombinant C1QL3-2HA made in HEK-293T cells (1 µl of conditioned media). Asterisk (*) denotes a non-specific band seen in both genotypes. Right: Blue Native PAGE of C1QL3-2HA purified from brain to reveal endogenous oligomeric formations. **D)** Littermates were tested in a standard 15-minute open-field behavior assay. Each dot is 1 mouse. **(E)** *C1ql3* qRT-PCR on the indicated homozygous mouse lines. *Gapdh* levels used for normalization. Data are mean ± SEM. Sample size indicated in each bar. 1-way ANOVA with post-hoc Bonferroni test; ** *p*=0.006.

We previously found that C1QL3 made in HEK293T cells forms high molecular weight oligomers, mostly hexamers and 12-mers (Sticco et al., 2021); however, whether endogenous C1QL3 has similar multimerization is unknown. To resolve this, we affinity-purified C1QL3-2HA from *C1ql3*^2HA^ mouse brains and performed Blue Native PAGE side-by-side with recombinant C1QL3-2HA from HEK293T cells. We found that the detectable bands after Western blot were similar (Figure 2C), revealing that endogenous C1QL3-2HA also multimerizes into approximately equal populations of hexamers and 12-mers.

We then used the open-field assay to test whether the 2HA insertion caused a behavioral phenotype in *C1ql3*^2HA^ mice compared to *C1ql3^+/+^* littermates. We previously demonstrated that *C1ql3^-/-^* (KO) and *C1ql3^+/-^* mice exhibited hyperactivity (Martinelli et al., 2016). No such phenotype was observed in *C1ql3*^2HA^ mice, suggesting that the 2HA KI allele causes no detriment to C1QL3 function (Figure 2D). Lastly, using qRT-PCR, we detected similar mRNA expression levels after harvesting total RNA from *C1ql3*^2HA^ and *C1ql3^+/+^* mouse brain cerebral cortices (Figure 2E), suggesting that the allele is not hypomorphic.

#### Comparison of C1ql3*^flox-mVenus^* and C1ql3*^2HA^*mice

We previously created a *C1ql3* allele with an IRES-mVenus (*C1ql3^flox-mVenus^*) knocked into the 3’ UTR (Martinelli et al., 2016), which exhibited a similar expression pattern to that observed with *in situ* hybridization (Iijima et al., 2010). Note that the cytoplasmic mVenus is not a fusion protein and fills cells expressing the bicistronic mRNA. To compare protein-level and mRNA reporter expression, we generated compound heterozygous mice (*C1ql3^flox-mVenus/2HA^*) (Fig. 3A). In many forebrain sections, including the cortex, cortical subplate, hippocampus, thalamus, and hypothalamus, we observed that nearly every cell expressing mVenus also showed HA-IR (Fig. 3B-D). However, we also observed numerous cell clusters that exhibited HA-IR but lacked detectable mVenus signal, *e.g.,* in the deep cerebellar nuclei (Fig. 3E), among other regions. This was reminiscent of our recent reports on *C1ql1* alleles, in which we found that the *C1ql1^flox-mVenus^* allele was hypomorphic with reduced mRNA levels, and the mVenus reporter did not always reveal a true *C1ql1*-expressing cell (Altunay et al., 2025; Biswas et al., 2021; Cheung et al., 2024). After testing *C1ql3* transcript levels in each genotype, we similarly observed that the *C1ql3^flox-mVenus^* allele was also hypomorphic (Figure 2E). We conclude that the HA tag is more likely to be a reliable reporter and *C1ql3^HA/HA^* mice were used for all subsequent experiments.

**Figure 3:**
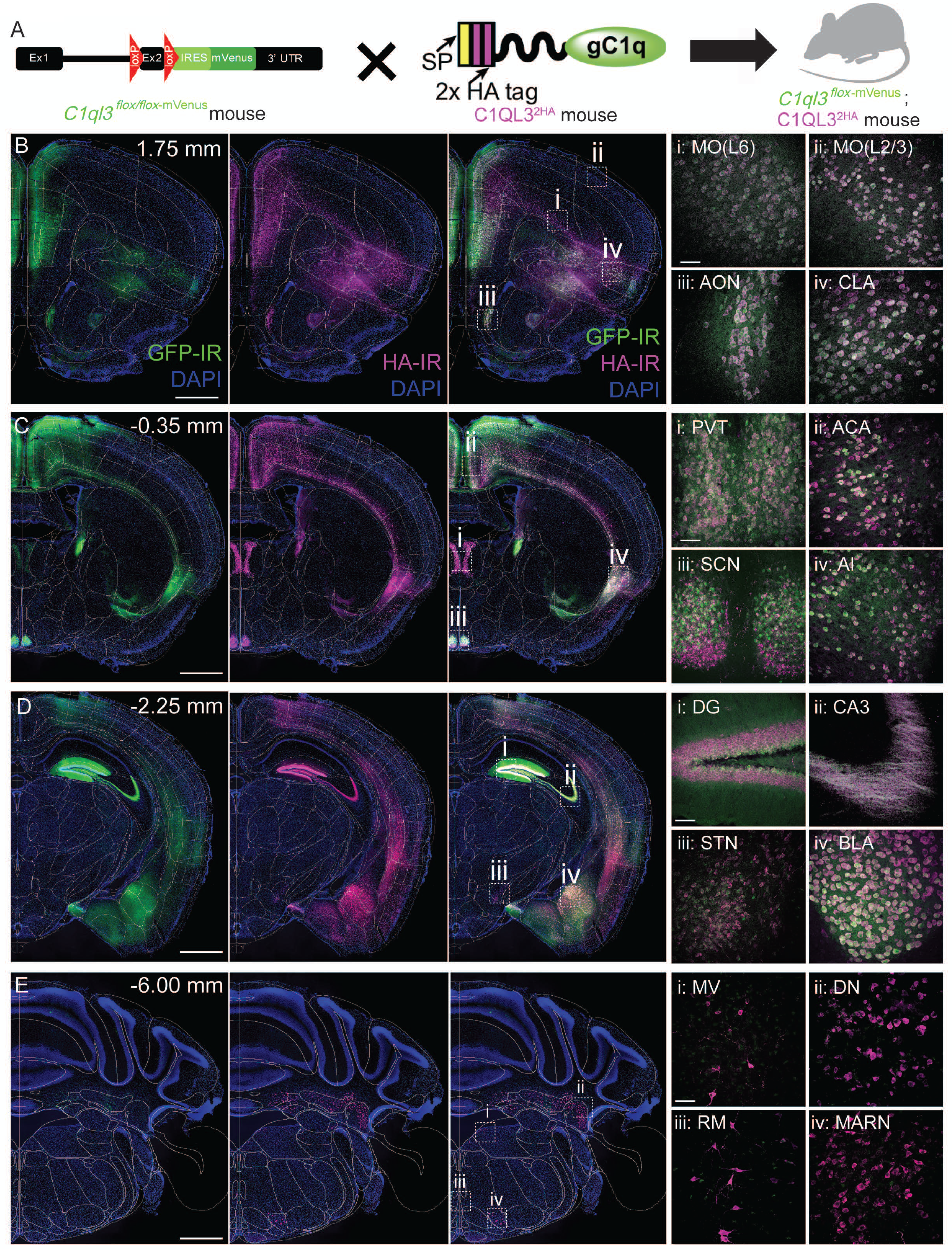
Comparison of novel *C1ql3^2HA^* mouse to the *C1ql3^-flox-^*^mVenus^ genetic reporter. **A)** Breeding of the *C1ql3-*mVenus mouse to the *C1ql3^2HA^* mouse to generate a compound heterozygote. **B)** Expression of *C1ql3-*mVenus (GFP-IR) and C1QL3-2HA (HA-IR) at Bregma level 1.75 mm (10X), highlighting co-expression in the: i) Primary motor cortex (MOp) layers 2/3, ii) MOp layer 6, iii) anterior olfactory nucleus (AON), and iv) claustrum (CLA; i-iv: 40X confocal). **C)** Expression at Bregma level -0.35 mm (10X), highlighting co-expression in the i) paraventricular thalamus (PVT), ii) anterior cingulate area (ACA), iii) suprachiasmatic nucleus (SCN), and iv) insular cortex (IC; i-iv: 40X confocal). **D)** Expression at Bregma level -2.25 mm (10X), highlighting co-expression in the i) dentate gyrus (DG), ii) CA3 of the hippocampus, iii) subthalamic nucleus (STN), and iv) basolateral amygdala (BLA; i-iv: 40X confocal). **E)** Expression at Bregma level -6.00 mm (10X), highlighting dim or lacking co-expression in the i) medial vestibular nucleus (MVN), ii) dentate nucleus (DN), iii) raphe magnus (RM), and iv) magnocellular reticular nucleus (MARN; i-iv: 40X confocal). (Scale bars= 1 mm for 10X, 50 µm for 40X)

### Brain-wide mapping of C1QL3-2HA using tissue clearing and light-sheet microscopy

To comprehensively map C1QL3 expression and potentially identify new or previously overlooked anatomical regions and cell populations that express C1QL3, we conducted whole-brain immunolabeling and light-sheet imaging of six adult *C1ql3*^2HA^ mice (∼3 months old, 3 male, 3 female) with *LifeCanvas Technologies*. Brains were cleared using SHIELD/Clear+ processing, labeled with anti-HA antibodies, and registered to the Allen Mouse Brain Atlas (mouse.brain-map.org and atlas.brain-map.org). Across all animals, regardless of sex, we observed consistent and robust C1QL3 expression patterns. Negative control immunohistochemistry (IHC) experiments for HA in wild type C57BL/6J mice confirmed that the HA-IR observed in *C1ql3*^2HA^ mice was not artifactual (Fig. 4A-E). Guided by publicly available coronal sections of *C1ql3* expression from the Allen Brain Cell Atlas MERFISH database (knowledge.brain-map.org) (Allen Institute & Zeng, 2023; Yao et al., 2023), we compared C1QL3-2HA expression to *C1ql3* mRNA expression in closely corresponding sections (Fig. 5). Although the coronal sections did not always perfectly align with our light-sheet imaged brains, the *C1ql3* mRNA expression patterns strikingly resembled HA-IR patterns in the light-sheet dataset (Fig. 5). The fluorescent signals observed along the edge of the lateral ventricles in the light-sheet images (Fig. 5C, D) were likely artifacts of tissue processing, as they were absent in HA-immunostained cryosections from both wild type and *C1ql3*^2HA^ mice (Fig. 4F). Brain regions with particularly high densities were noted, quantified as average density (10^3^ cells/mm^3^) per subregion across 6 individual mice (Fig. 6).

**Figure 4:**
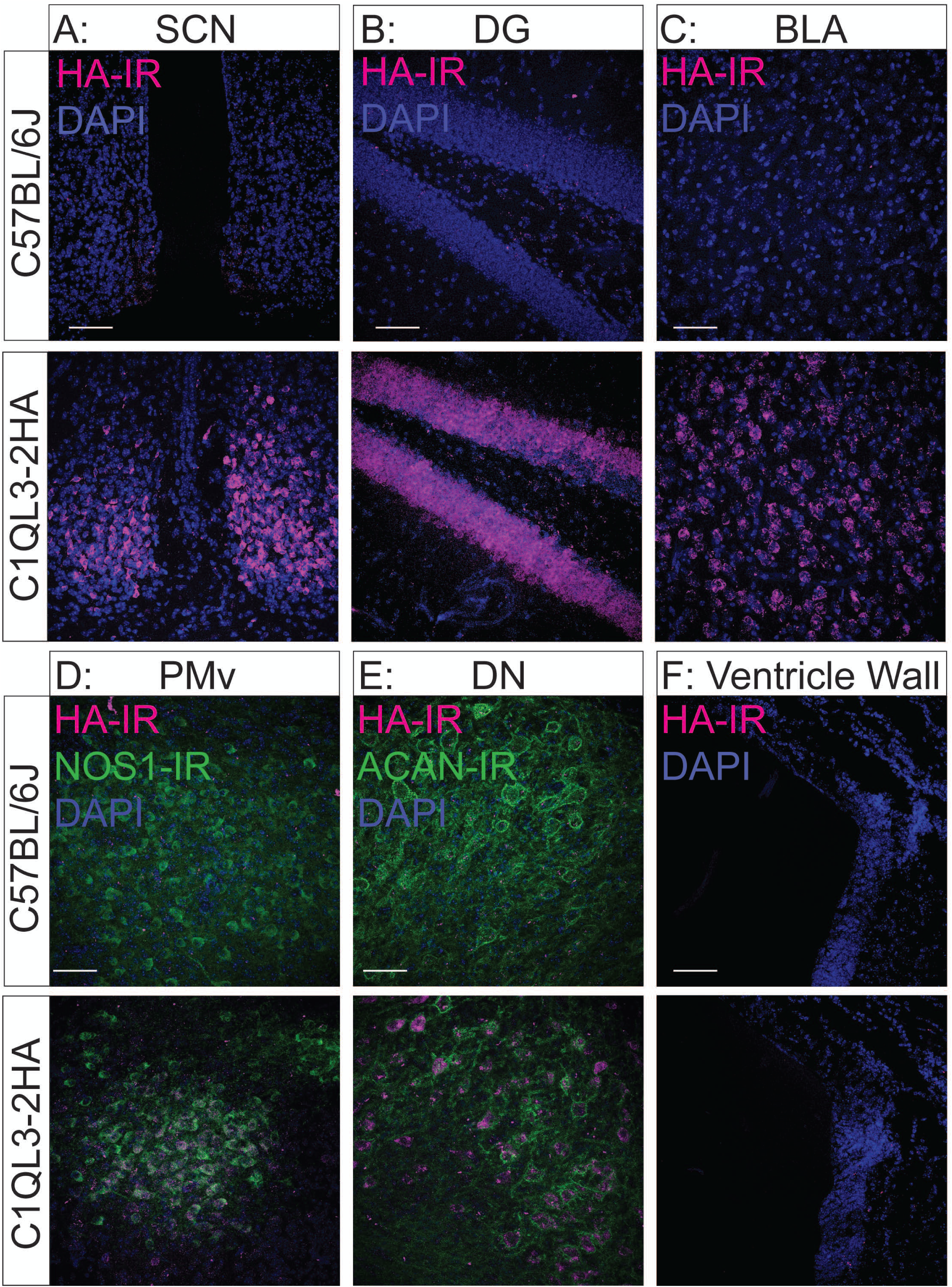
Negative control for anti-HA staining. Comparison of HA immunohistochemistry in a C57BL/6J mouse and age-matched adult C1QL3^2HA^ mouse, demonstrating the specificity of HA-IR. **A)** Suprachiasmatic nucleus (SCN). **B)** Dentate gyrus (DG). **C)** Basolateral amygdala (BLA). **D)** Ventral premammillary nucleus (PMv) counterstained with anti-nitric oxide synthase 1 (NOS1). **E)** Dentate nucleus (DN) of the deep cerebellar nuclei counterstained with anti-aggrecan (ACAN). **F)** Lateral ventricular wall at approximate Bregma level -0.5 mm. (All scale bars=50 µm)

**Figure 5:**
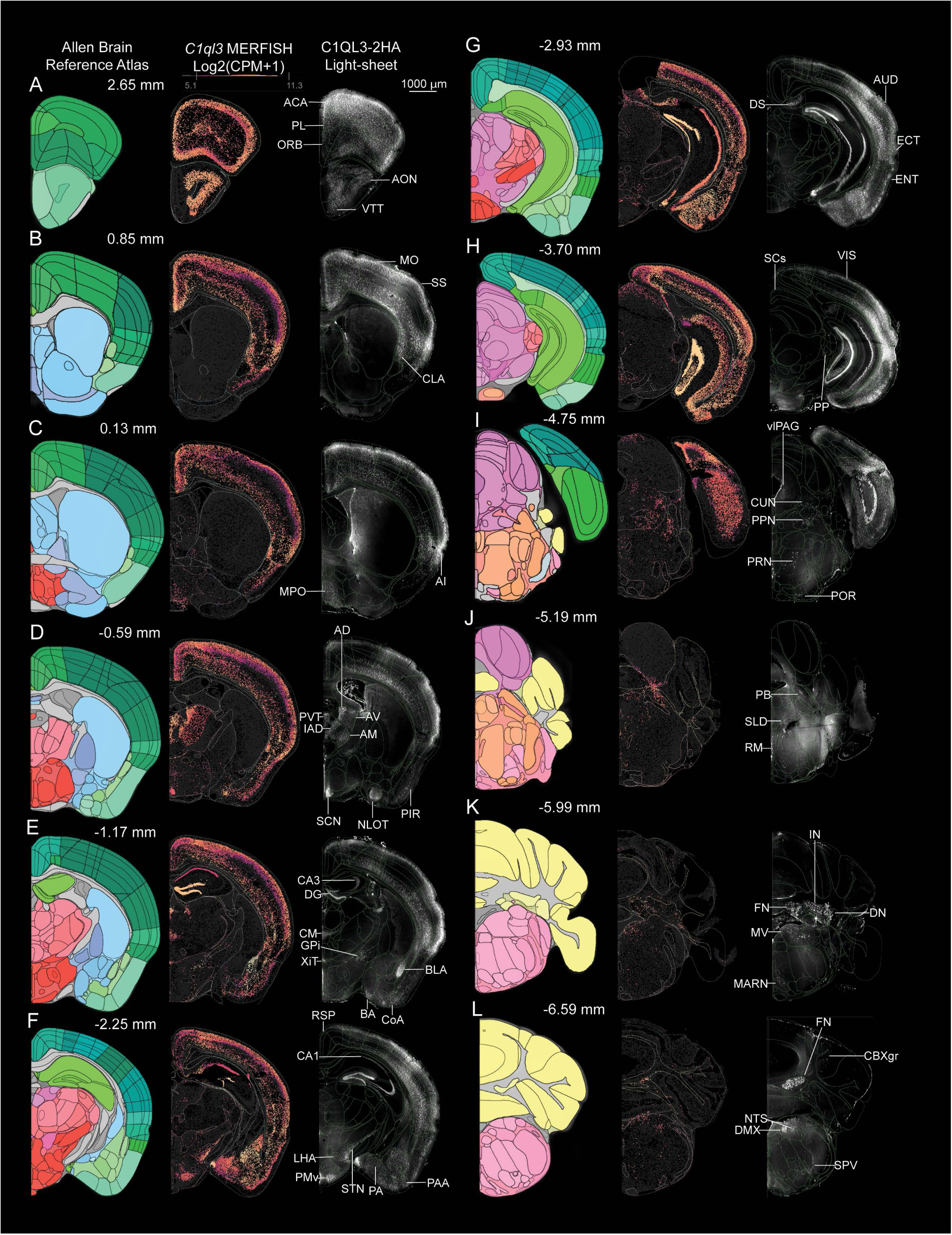
Brain-wide expression of *C1ql3* mRNA and HA-IR in C1QL3-2HA mice. Side-by-side comparison of select coronal sections from the Allen Mouse Brain Reference Atlas (left column), *C1ql3* mRNA expression from the Allen Mouse Brain MERFISH Atlas (middle column), and corresponding C1QL3-2HA-IR in light-sheet imaging, along the anteroposterior axis at the indicated approximate Bregma levels. **A)** 2.65 mm, highlighting the anterior cingulate area (ACA), prelimbic (PL) and orbital (OC) cortices, anterior olfactory nucleus (AON), and ventral taenia tecta (VTT). **B)** 0.85 mm, highlighting the motor (MO) and somatosensory (SS) cortices and the claustrum (CLA). **C)** 0.13 mm, highlighting the agranular insula (AI) and medial preoptic area of the hypothalamus (MPO). **D)** -0.59 mm, highlighting the anterodorsal (AD), paraventricular (PVT), interanterodorsal (IAD), anteroventral (AV) and anteromedial (AM) thalamic nuclei, suprachiasmatic nucleus (SCN), nucleus of the lateral olfactory tract (NLOT), and piriform cortex (PIR). **E)** -1.17 mm, highlighting the dentate gyrus (DG) and CA3 region of the hippocampus, centromedial (CM) and xiphoid (XiT) thalamic nuclei, basolateral (BLA) and cortical (CoA) amygdalar nuclei, and bed nucleus of the accessory olfactory tract (BA). **F)** -2.25 mm, highlighting the CA1 region of the hippocampus, ventral premammillary nucleus (PMv), subthalamic nucleus (STN), posterior amygdala (PA) and piriform-amygdalar-area (PAA). **G)** -2.93 mm, highlighting the dorsal subiculum (DS) and auditory (AUD), ectorhinal (ECT), and entorhinal (ENT) cortices. **H)** -3.70 mm, highlighting the superior colliculus (SC), visual cortex (VIS) and peripeduncular thalamus (PP). **I)** -4.75mm, highlighting the ventrolateral periaqueductal grey (vlPAG), cuneiform nucleus (CUN), pedunculopontine nucleus (PPN), pontine reticular nucleus (PRN), and periolivary region (POR). **J)** -5.19 mm, highlighting the parabrachial nucleus (PB), sublaterodorsal tegmentum (SLD) and raphe magnus (RM). **K)** -5.99 mm, highlighting the fastigial (FN), interposed (IN), and dentate (DN) deep cerebellar nuclei, the medial vestibular nucleus (MV) and magnocellular reticular nucleus of the medulla (MARN). **L)** -6.59 mm, highlighting the granule cell layer of the cerebellar cortex (CBXgr), nucleus of the solitary tract (NTS), dorsal nucleus of the vagus nerve (DMX), and spinal trigeminal nucleus (SPV).

**Figure 6:**
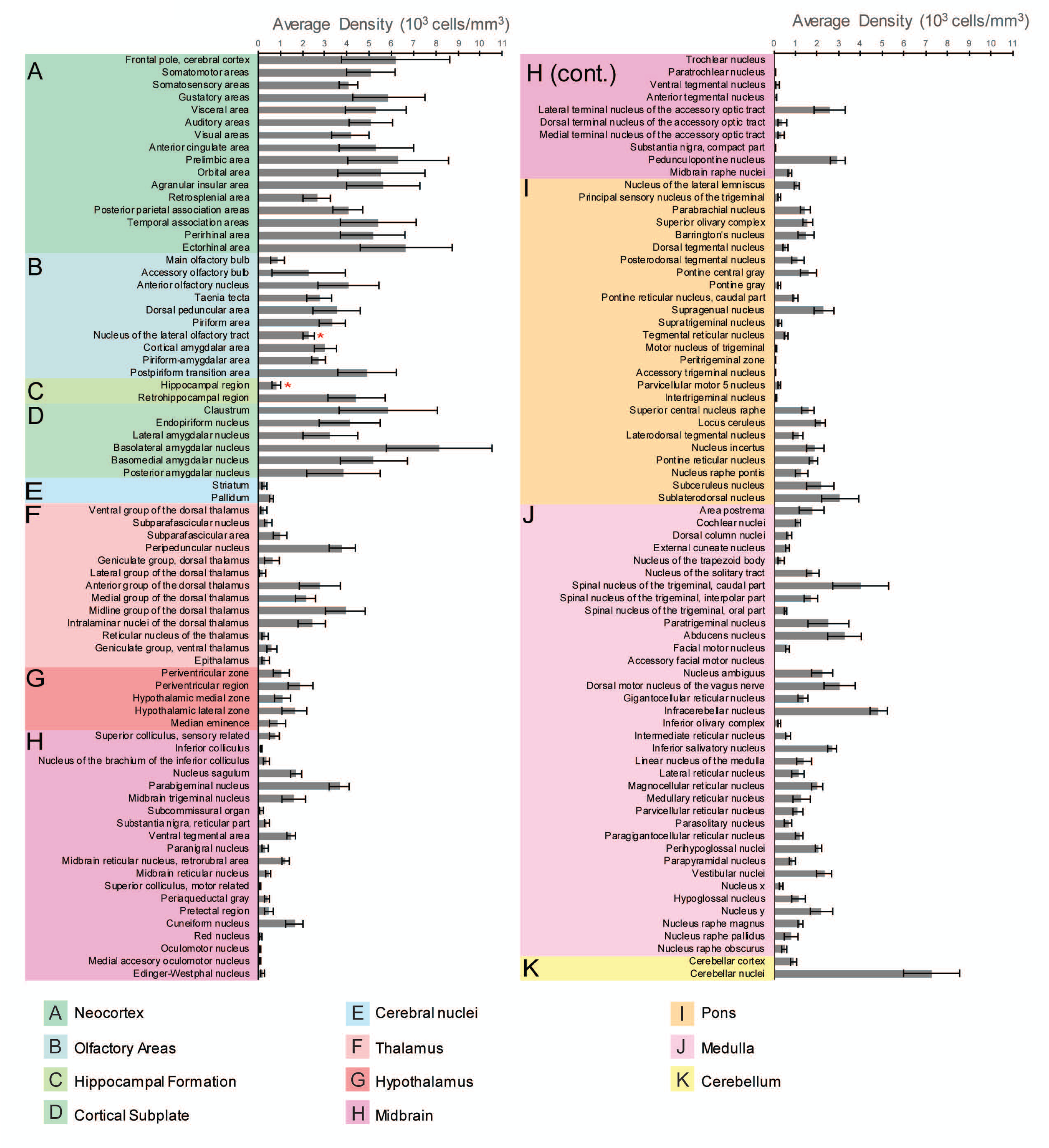
Cell density quantification of C1QL3-2HA light-sheet data. Cell density (as average density of cells, 10^3^ cells/mm^3^) of C1QL3-2HA light-sheet data (n=6 mice) across anatomical subregions as defined by the Allen Mouse Brain Atlas. **A)** Neocortex. **B)** Olfactory areas. **C)** Hippocampal formation. **D)** Cortical subplate. **E)** Cerebral nuclei. **F)** Thalamus. **G)** Hypothalamus. **H)** Midbrain. **I)** Pons. **J)** Medulla. **K)** Cerebellum. (* indicates regions in which cells are too dense to accurately quantify).

We observed robust and widespread C1QL3-2HA expression throughout the forebrain, midbrain, hindbrain, and cerebellum in the light-sheet/MERFISH data (Fig. 5) and light-sheet quantification (Fig. 6) as described in more detail in the following sections. Notably, C1QL3-2HA-positive cells are highly enriched in layers 2/3 and 6 of the neocortex, the claustrum, the anterior olfactory nucleus, deep cerebellar nuclei, and several thalamic and hypothalamic nuclei. Previously underappreciated expression was identified in the dorsal subiculum, superior colliculus, and brain regions enriched in monoaminergic neurons, including the VTA, raphe nuclei, and LC. Region-specific cell density quantification revealed that C1QL3-2HA is expressed in both excitatory and inhibitory neurons, with laminar patterns in cortex and highly restricted subnuclear distributions in subcortical regions (Fig. 6).

### C1QL3-2HA expression in the telencephalon

The newly generated *C1ql3*^2HA^ KI mouse revealed a highly structured expression pattern of C1QL3-2HA across the telencephalon. C1QL3-2HA positive (C1QL3-2HA+) cells were densely distributed across all subdivisions of the neocortex (Fig. 6A). The medial prefrontal cortex (mPFC), including the prelimbic (PL), orbital (ORB), and anterior cingulate (ACA) cortices, showed robust and continuous expression (Fig. 5A), which gradually tapered in the posterior retrosplenial cortex (RSP, Fig. 5F).

The layer-specific organization of C1QL3-2HA+ neurons became particularly pronounced in the motor (MO), somatosensory (SS), auditory (AUD), and visual cortices (Fig. 5B, G, H), indicating a laminar precision of C1QL3 expression that may underlie distinct circuit roles. This patterning was notably absent in transitional zones, such as the insular cortex (AI, Fig. 5C) and the ectorhinal cortex (ECT, Fig. 5G), regions that bridge neocortical and allocortical territories. Yet, both were packed with HA positive (HA+) cells, suggesting specialized roles at cortical boundaries. Within the allocortex itself, the expression pattern changed: in the anterior piriform cortex (PIR, Fig. 5D), C1QL3-2HA+ cells were confined to a narrow layer, while the entorhinal cortex (ENT, Fig. 5G and quantified in Fig. 6C) contained HA+ neurons across multiple layers. We observed a pattern of C1QL3-2HA expression in olfactory forebrain structures (Fig. 6B), where labeling was highly selective. While absent from most of the main olfactory bulb, a dense population of HA+ neurons emerged in the anterior olfactory nucleus (AON, Fig. 5A) and the ventral taenia tecta (VTT, Fig. 5A). Most strikingly, the nucleus of the lateral olfactory tract (NLOT, Fig. 5D) exhibited such a dense HA+ cell cluster that the automated cell counting algorithm could not fully resolve it. High expression in the bed nucleus of the accessory olfactory tract (BA, Fig. 5E) and amygdalopiriform area (PAA, Fig. 5F) adds to the emerging view that C1QL3 may be a central player in olfactory-limbic integration (Wang et al., 2020).

In the hippocampus, the *C1ql3*^2HA^ mouse revealed a uniform distribution of C1QL3-2HA+ granule cells in the dentate gyrus (DG, Fig. 5E), consistent with prior studies (Fig. 6C) (Iijima et al., 2010; Martinelli et al., 2016). Low-density HA+ neurons were seen in the CA1 subfield (Fig. 5F), and dense mossy-fiber axonal labeling is visible in CA3 (Fig. 5E and Fig. 3Dii), reflecting C1QL3 trafficking within axon terminals. Although synaptic signal detection was limited without antigen retrieval, these results hint at a strong presynaptic presence. The retrohippocampal area showed dense HA+ expression in the dorsal subiculum (DS, Fig. 5G) and widespread labeling in the entorhinal cortex.

In cortical subplate-derived regions, we observed dense C1QL3-2HA signal in both the claustrum (CLA, Fig. 5B) and the basolateral amygdala (BLA, Fig. 5E), the latter previously implicated in C1QL3 function (Martinelli et al., 2016). Additional amygdaloid nuclei, including the cortical (CoA, Fig. 5E) and posterior amygdala (PA, Fig. 5F), also exhibited discrete C1QL3-2HA expression (Fig. 6D), suggesting a broader role in emotional and sensory processing networks.

In contrast, C1QL3-expressing soma were absent from the striatum and pallidum (Fig. 6E), sharply delineating regions of enrichment and relative absence. Note that without antigen retrieval, we would not detect any pre-synaptic C1QL3 protein from cortico-striatal projections. A notable exception appeared in the internal segment of the globus pallidus (GPi), or entopeduncular nucleus, where a small cluster of dim HA+ cells was visible (Fig. 5E). This highly selective expression pattern invites further investigation with regard to potential modulatory roles of C1QL3 beyond classical excitatory circuits.

### C1QL3-2HA expression in the diencephalon

The *C1ql3*^2HA^ mouse revealed a sharply demarcated pattern of C1QL3 expression across the diencephalon, much of which was not previously appreciated or detected in prior studies. This feature was evident in the thalamus, where C1QL3-2HA+ cells were confined to distinct subnuclei and absent from surrounding regions (Fig. 6F). This exclusivity was particularly striking in the anterior and midline thalamic nuclei which exhibit the highest densities of HA+ cells, pointing to a potential role of C1QL3 in circuits involved in attention, arousal, and limbic processing.

Among the most enriched areas in the diencephalon were the paraventricular thalamus (PVT, Fig. 5D), in agreement with prior work (Iijima et al., 2010; Martinelli et al., 2016), as well as the centromedial (CM, Fig. 5E), interanterodorsal (IAD, Fig. 5D), and anterodorsal (AD, Fig. 5D) nuclei. These regions anchor thalamo-limbic loops (Colavito et al., 2015; Kooiker et al., 2021; Vertes & Hoover, 2008), suggesting that C1QL3 could be a key molecular player in modulating internal state and behavioral flexibility. Even within closely neighboring nuclei, the precision of expression was remarkable: the anteroventral (AV, Fig. 5D) and anteromedial (AM, Fig. 5D) thalamic nuclei showed dimmer but still discernible HA+ signals, while the vast majority of the posterior thalamus was devoid of C1QL3-2HA+ soma. The only exception, the peripeduncular nucleus (PP, Fig. 5H), contained a small cluster of HA+ cells, standing out as a rare posterior thalamic site of expression.

In the hypothalamus, C1QL3-2HA expression was again sparse at the global level but revealed key hotspots of expression that may have important functional implications (Fig. 6G). The suprachiasmatic nucleus (SCN, Fig. 5D) was a clear standout, showing intense and localized C1QL3-2HA labeling consistent with our previous report (Chew et al., 2017). Beyond the SCN, additional pockets of expression emerged that had previously received little attention in the context of C1QL3 biology. Notably, the medial preoptic area (MPO, Fig. 5C) contained scattered HA+ neurons, suggesting a potential role in reproductive or homeostatic regulation (Rothhaas & Chung, 2021). In the lateral hypothalamic area (LHA, Fig. 5F), posterior hypothalamus (PH), the ventral premammillary nucleus (PMv, Fig. 5F) and the subthalamic nucleus (STN, Fig. 5F) exhibited distinct enrichment of HA+ cells, suggesting roles for C1QL3 in modulating hypothalamic outputs related to motivation, feeding, sleep-wake, locomotion and affective states (Bonnavion et al., 2016; Bonnevie & Zaghloul, 2019; Cavalcante et al., 2025; Fong et al., 2023).

### C1QL3-2HA expression in the midbrain, brainstem and cerebellum

Moving posteriorly, the C1QL3^2HA^ mouse continued to reveal previously undetected or unappreciated C1QL3-expressing subcortical nuclei. In the midbrain (Fig. 6H), several subnuclei displayed C1QL3-2HA expression patterns. A particularly dense cluster of HA+ neurons was identified in the ventrolateral periaqueductal gray (vlPAG, Fig. 5I), a key node in pain modulation and behavioral responses to threat (Falkner et al., 2020; Li et al., 2018). The lateroposterior midbrain also stood out: the pedunculopontine nucleus (PPN) and cuneiform nucleus (CUN, Fig. 5I) were both populated with HA+ cells, suggesting a role in locomotor control and arousal (Lau et al., 2015; Mena-Segovia & Bolam, 2017a; Ryczko & Dubuc, 2013). Notably, we also detected HA+ neurons in the dorsal sensory layers of the superior colliculus (SCs, Fig. 5H), in contrast to the absence of labeling in the deeper collicular subdivisions. This intriguing pattern suggests sensory-specific involvement in visual-motor integration (Liu et al., 2022).

In the pons (Fig. 6I), C1QL3 expression also exhibited distinct anatomical expression patterns. A well-defined band of HA+ neurons appeared in the pontine reticular nucleus (PRN, Fig. 5I). Additional HA+ populations were identified in the parabrachial nucleus (PB, Fig. 5J), the sublaterodorsal tegmentum (SLD, Fig. 5J), and the periolivary region (POR, Fig. 5I), all of which are functionally implicated in autonomic regulation, sleep, and auditory processing (Chiang et al., 2019; Torontali et al., 2019; Vetter, 2015). Scattered HA+ cells were also observed throughout the dorsal pons, indicating a broader distribution than previously appreciated.

The medulla (Fig. 6J) displayed several robust clusters of C1QL3-2HA+ cells in functionally diverse areas. Prominent among these were the large HA+ neurons of the raphe magnus (RM, Fig. 5J), suggesting a potential role for C1QL3 in serotonergic signaling. Other divisions of the midbrain raphe contained fewer and dimmer HA+ cells, indicating a more selective involvement. The medial vestibular nucleus (MV, Fig. 5K) showed widespread labeling, alongside scattered expression across other vestibular subdivisions, suggesting a possible role for C1QL3 in balance and spatial orientation (Khan & Chang, 2013). The magnocellular reticular nucleus (MARN, Fig. 5K) exhibited HA+ cells, consistent with a potential role in motor pattern generation (Brownstone & Chopek, 2018). A band of HA+ neurons was observed in the dorsal portion of the nucleus of the solitary tract (NTS, Fig. 5L), a central hub for visceral sensory integration (Huang et al., 2025; Zoccal et al., 2014). Scattered expression was also seen in the spinal trigeminal nucleus (SPV, Fig. 5L), which processes somatosensory input from the head and face (Henssen et al., 2016). Strikingly, a dense cluster of HA+ cells sharply delineated the dorsal motor nucleus of the vagus nerve (DMX, Fig. 5L), positioning C1QL3 as a candidate regulator of parasympathetic outflow (Strain et al., 2024).

Finally, the cerebellum (Fig. 6K) showed robust C1QL3-2HA expression in all three deep cerebellar nuclei: dentate (DN), interposed (IN), and fastigial (FN, Fig. 5K-L). Each of these nuclei serves as a major output center for the motor and cognitive functions of the cerebellum (Okada et al., 2022). In the cerebellar cortex, sparse HA+ cells were observed within the granule cell layer (CBXgr, Fig. 5L), suggesting limited but potentially important cerebellar cortical involvement.

These data reveal new areas of expression that implicate C1QL3 in a wide range of functions, including sensory gating, motor execution, autonomic control, and arousal regulation, providing a model for the functional dissection of C1QL3 in integrative neural systems. The following sections expand on select brain regions and cell populations enriched in C1QL3-2HA.

### Laminarization of C1QL3-2HA expression across cortical regions

The consistent layer-specific expression of C1QL3-2HA across neocortical subdivisions prompted a deeper investigation into its cortical laminar organization. Quantitative line-scan analysis of fluorescence intensity across the cortical depth revealed a striking and reproducible laminar profile. In the primary motor cortex (MO), we found a near-complete absence of signal in layer 1, aside from minor edge artifacts, followed by a sharp peak in superficial layers 2/3, consistent with a dense band of HA+ somata in this region (Fig. 7A–B). Fluorescence intensity markedly declined across layers 4 and 5, before showing a modest increase in layer 6, where weaker, yet clearly labeled, HA+ neurons were consistently detected.

**Figure 7:**
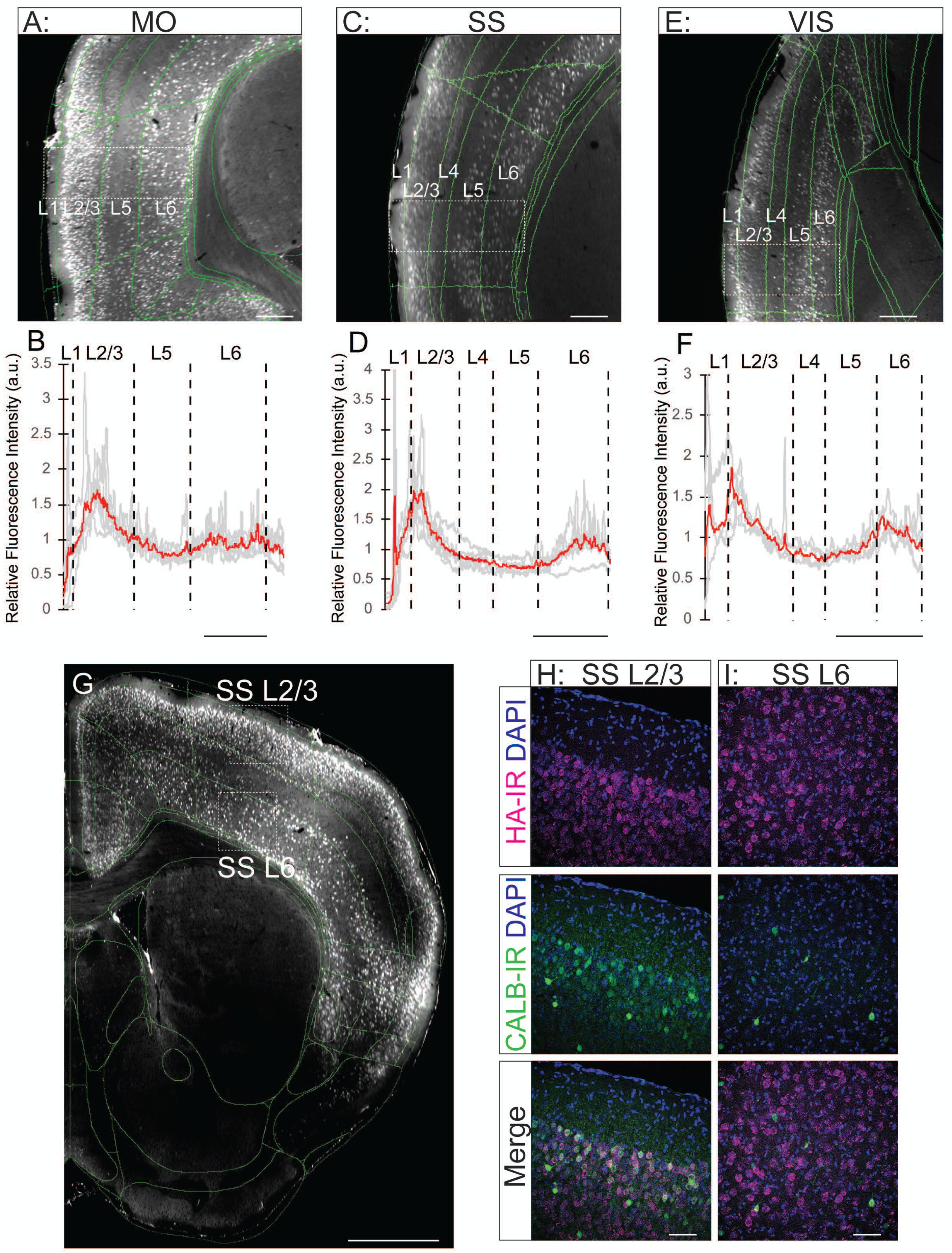
Laminarization of C1QL3-2HA expression in select regions of the neocortex. **A)** HA-IR visualized by light-sheet imaging in the layers of the motor cortex (MO) (image rotated to match layer quantification in B, scale bar=100 µm). **B)** Quantification of fluorescence intensity averaged across 5 close-proximity line scans from layer 1 to layer 6 of the MO. (scale bar=100 µm) **C)** HA-IR visualized by light-sheet imaging in the layers of the somatosensory (SS) cortex (image rotated to match layer quantification in D scale bar=100 µm). **D)** Quantification of fluorescence intensity averaged across 5 close-proximity line scans from layer 1 to layer 6 of the primary SS (scale bar=100 µm). **E)** HA-IR visualized by light-sheet imaging in the layers of the visual cortex (VIS) (image rotated to match layer quantification in F, scale bar=100 µm). **F)** Quantification of fluorescence intensity averaged across 5 close-proximity line scans from layer 1 to layer 6 of the VIS (scale bar=100 µm). **G)** Representative light-sheet microscopy image of HA-IR in the neocortex at bregma level 1.53 mm. (Scale bar=1 mm) **H)** Co-localization of HA-IR with calbindin (CALB)-IR in L2/3 of the primary MO of the C1QL3^2HA^ mouse. **I)** Lack of co-localization of HA-IR with CALB-IR in L6 of the primary MO of the C1QL3^2HA^ mouse. (Scale bar=50 µm)

This laminar expression pattern was conserved across other neocortical regions, including the somatosensory (SS) and visual cortices (VIS) (Fig. 7C–F), underscoring a potentially shared structural principle of C1QL3 organization in neocortical circuits. To further define cell type-specificity, we performed dual IHC for HA and calbindin (D28K) in the SS (Fig. 7G). In layers 2/3, where calbindin marks some pyramidal neurons (DeFelipe, 1997), we observed strong colocalization with HA, indicating that C1QL3-2HA is robustly expressed in excitatory projection neurons (Fig. 7H). In contrast, in layer 6, where calbindin marks a subset of interneurons, there was no overlap with HA (Fig. 7I), supporting the conclusion that C1QL3 expression in the cortex is restricted to excitatory neuronal populations and avoids GABAergic interneurons.

### C1QL3-2HA expression relative to specific subcortical cell populations

We leveraged our C1QL3^2HA^ mouse to gain cellular resolution into the identity of various C1QL3-expressing neurons across subcortical structures. In the thalamus, C1QL3-2HA expression resembles the known distribution of calretinin (Viena et al., 2021). Dual immunostaining confirmed colocalization in multiple nuclei, including the anterior dorsal thalamus (AD), where CALR-IR defined a medial subpopulation of HA+ cells (Fig. 8A–B). This relationship was even more pronounced in the paraventricular thalamus (PVT), where virtually all CALR+ neurons co-expressed HA (Fig. 8C), suggesting a strong functional link between C1QL3 and CALR-defined thalamic circuits.

**Figure 8:**
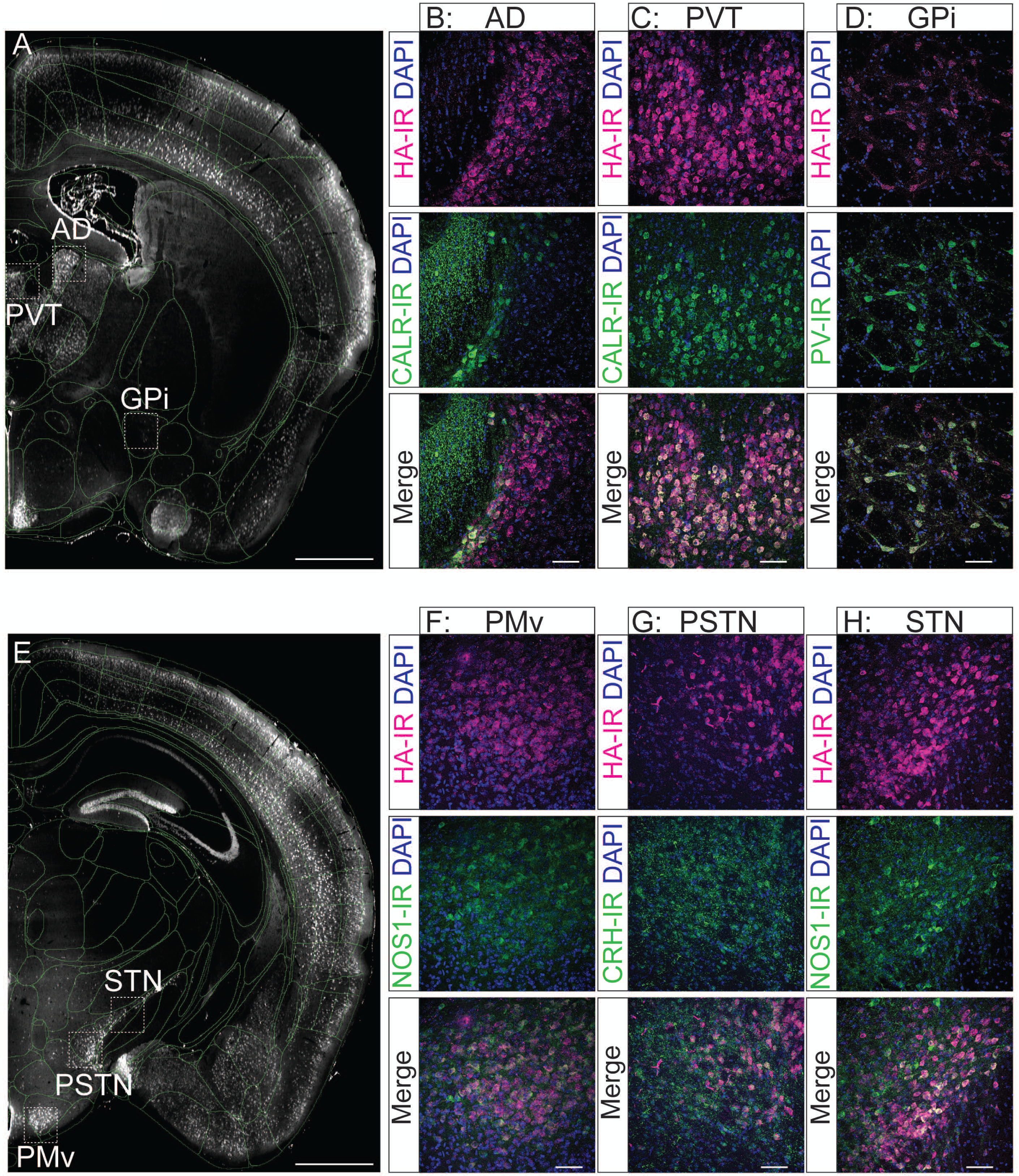
**Highlighted C1QL3-2HA+ cell populations in the pallidum and diencephalon.** Representative light-sheet microscopy images of **A)** HA-IR in the thalamus at Bregma level -0.59 mm, with **B)** co-localization of HA-IR with calretinin (CALR)-IR in the anterodorsal (AD) and **C)** paraventricular (PVT) thalamic nuclei, with **D)** co-localization of parvalbumin (PV)-IR and HA-IR in the globus pallidus internal part (GPi). Representative light-sheet microscopy image of **E)** HA-IR in the ventroposterior hypothalamus at bregma -2.45, with **F)** co-localization of HA-IR with nitric oxide synthase 1 (NOS1)-IR in the PMv, **G)** moderate co-localization of HA-IR with corticotropin-releasing hormone (CRH)-IR the parasubthalamic nucleus (PSTN), and **H)** moderate co-localization of HA-IR with NOS1-IR the STN. (Scale bars=1 mm for light-sheet images, 50 µm for confocal images)

Although the striatum and pallidum were largely devoid of C1QL3-2HA+ cells, we observed a small cluster of HA+ neurons in the internal globus pallidus (GPi/entopeduncular nucleus), a region known for dense parvalbumin (PV) expression (Miyamoto & Fukuda, 2015; Wallace et al., 2017). Immunostaining confirmed partial overlap of HA with PV-IR (Fig. 8D), indicating that C1QL3 is expressed in a distinct PV+ subpopulation, while additional HA+/PV− cells suggest potential molecular heterogeneity within the GPi, consistent with previous work (Miyamoto & Fukuda, 2015; Wallace et al., 2017).

In the ventroposterior hypothalamus and nearby regions of the diencephalon, C1QL3-2HA expression again revealed discrete molecular identities. Dual labeling showed moderate co-expression of HA and nitric oxide synthase 1 (NOS1) in both the ventral premammillary nucleus (PMv, Fig. 8F) and the subthalamic nucleus (STN, Fig. 8H), although HA-IR showed stronger somatic signal than NOS1 in the latter. In the parasubthalamic nucleus (PSTN), we observed moderate colocalization between HA and corticotropin-releasing hormone (CRH), consistent with expression in a defined PSTN neuronal population (Kim et al., 2022; Zhu et al., 2012), although the strong axonal labeling of CRH may have obscured some HA+ somata (Fig. 8G).

Progressing into the midbrain and brainstem, the spatial precision of C1QL3-2HA expression was also evident (Fig. 9A). In the raphe nuclei, we observed moderate co-labeling with tryptophan hydroxylase 2 (TPH2) in the dorsal raphe (DR, Fig. 9B) and the medullary raphe magnus (RM, Fig. 9C), confirming expression in some serotonergic neuron populations (Gutknecht et al., 2009; Okaty et al., 2015). Interestingly, in the pedunculopontine nucleus (PPN), a region enriched in cholinergic neurons (Mena-Segovia & Bolam, 2017b; Miyamoto & Fukuda, 2015), we detected robust C1QL3-2HA expression that did not overlap with choline acetyltransferase (ChAT)-IR, suggesting a distinct, non-cholinergic, but intermingled C1QL3+ neuronal population (Fig. 9D).

**Figure 9:**
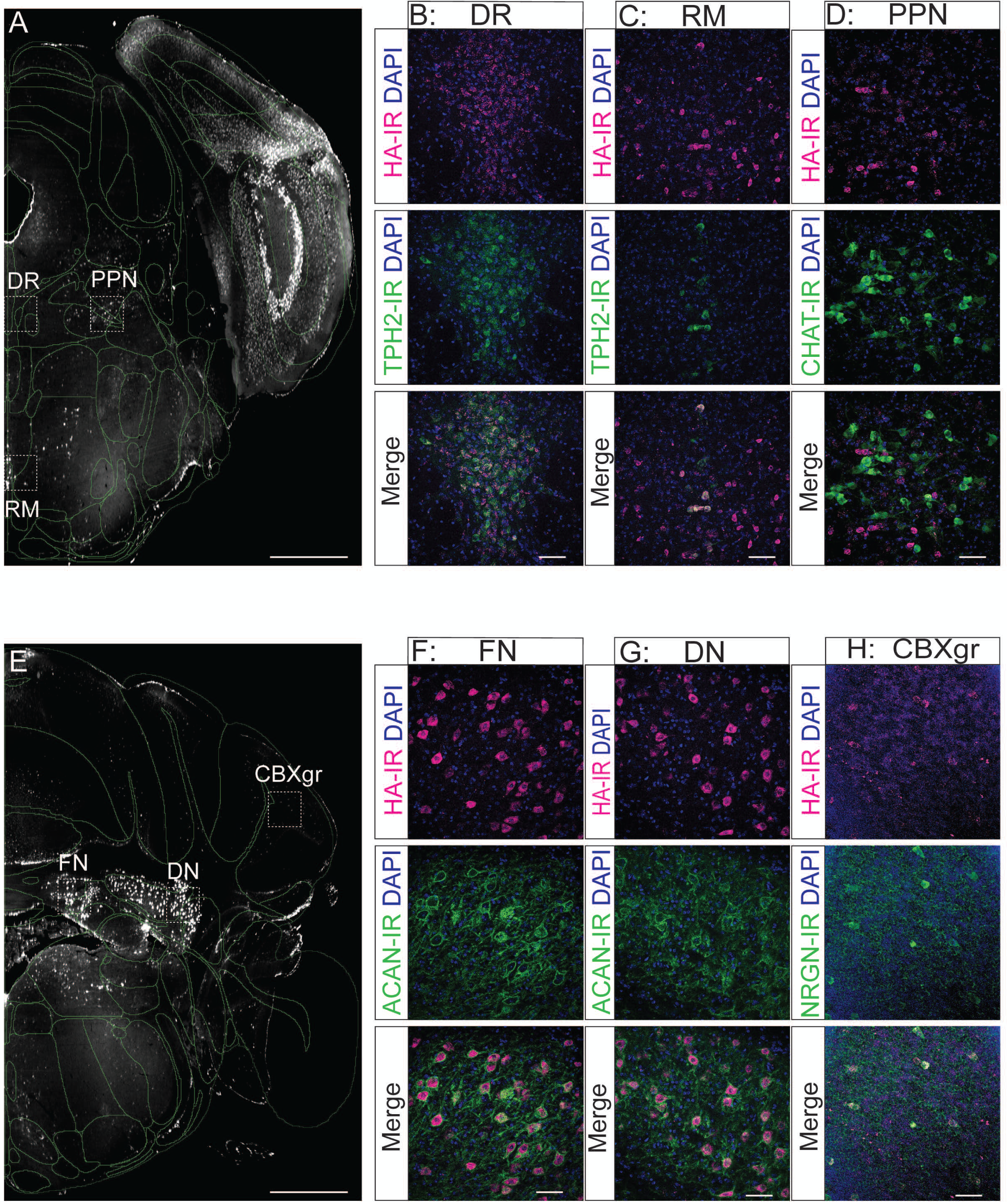
Highlighted C1QL3-2HA+ cell populations in the brainstem and cerebellum. Representative light-sheet microscopy images of **A)** HA-IR in the midbrain at Bregma level -5.20 mm with **B)** co-localization of HA-IR with tryptophan hydroxylase-2 (TPH2)-IR in the dorsal raphe (DR) and **C)** raphe magnus (RM) and **D)** Lack of co-localization of HA-IR with choline acetyl transferase (ChAT)-IR the pedunculopontine nucleus (PPN). Representative light-sheet microscopy images of **E)** HA-IR in the cerebellum at Bregma level -5.90 mm with **F)** co-localization of HA-IR with aggrecan (ACAN)-IR in fastigial cerebellar nucleus (FN) and **G)** dentate cerebellar nucleus (DN) and **H)** co-localization of HA-IR with neurogranin (NRGN)-IR marking Golgi interneurons in the granular cell-layer of the cerebellar cortex (CBXgr). (Scale bars=1 mm for light-sheet images, 50 µm for confocal images)

In the cerebellum, dual-labeling approaches further defined C1QL3-2HA-expressing cell types in both deep nuclei and cortical layers (Fig. 9E). Using aggrecan (ACAN) as a marker of deep cerebellar nuclear neurons (Matthews et al., 2002; Zaremba et al., 1990), we observed a high degree of colocalization between HA-IR and ACAN-IR across all deep cerebellar nuclei, including the fastigial (FN) and dentate subdivisions (DN) (Fig. 9F–G), validating the presence of C1QL3-2HA in projection neurons that shape cerebellar output. Additionally, in the granular layer of the cerebellar cortex, we observed partial colocalization of HA with neurogranin (NRGN), a marker of Golgi interneurons (Fig. 9H) (Simat et al., 2007), revealing a new cerebellar cell population potentially shaped by C1QL3.

Finally, we probed C1QL3-2HA expression in canonical catecholaminergic systems. In the ventral tegmental area (VTA), dual labeling with tyrosine hydroxylase (TH) revealed that C1QL3-2HA+ cells represent a small but distinct subset of dopaminergic neurons (Fig. 10A–B). In the locus coeruleus (LC), we also identified scattered HA+ cells; however, they did not colocalize with TH, suggesting that they are non-noradrenergic cells intermingled with LC noradrenergic neurons (Fig. 10C–D).

**Fig 10:**
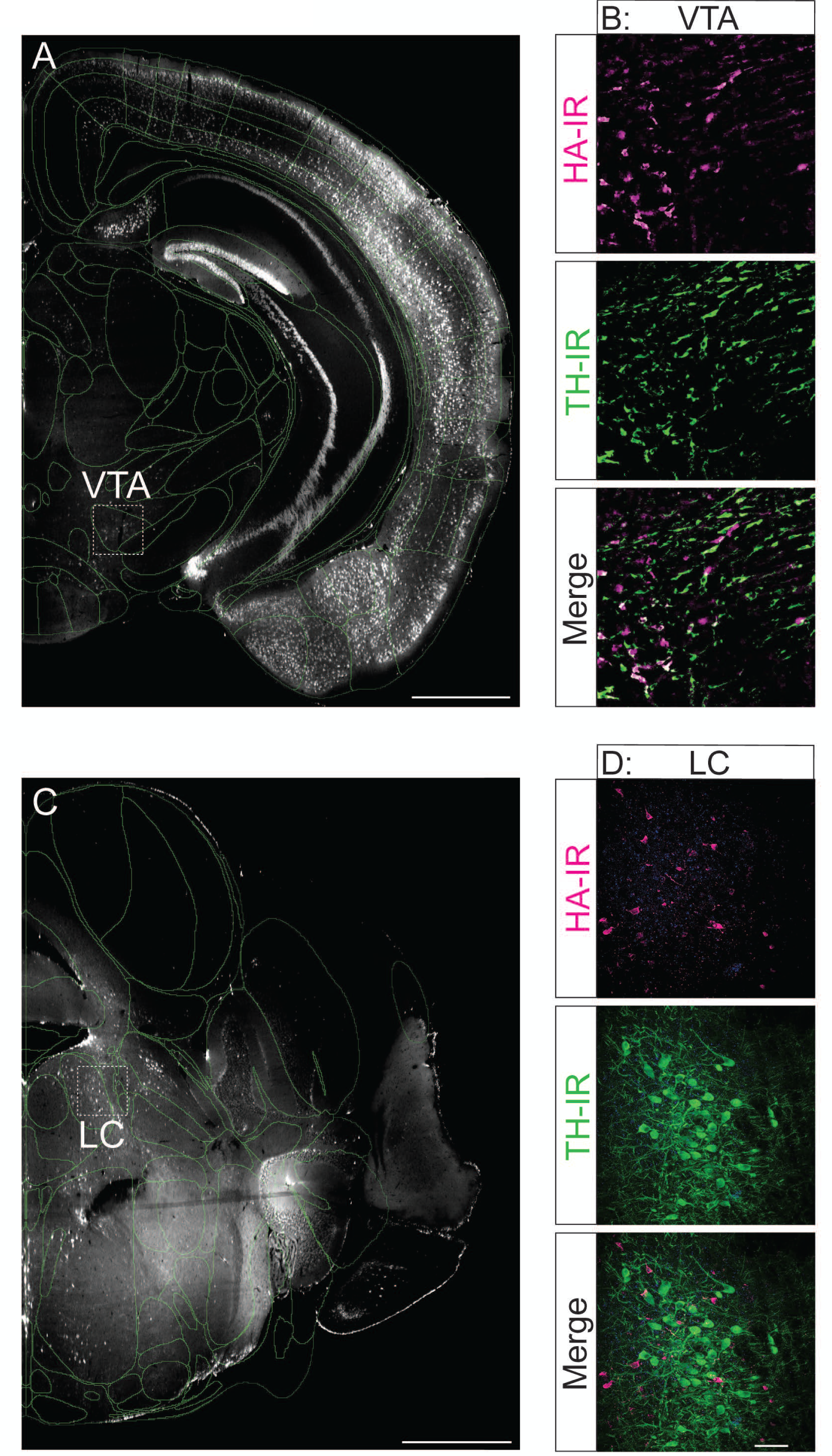
H**i**ghlighted **C1QL3-2HA expression in two catecholaminergic cell groups. A)** Dim HA+ cells visible in the ventral tegmental area (VTA) of the C1QL3^2HA^ mouse. **B)** HA-IR in the VTA which co-localizes with a subset tyrosine hydroxylase (TH)-IR dopaminergic neurons in the VTA (sectioned at 20 µm to improve signal to noise). **C)** HA+ cells visible in and around the locus coeruleus (LC). **D)** Lack of colocalization in the LC of HA-IR with TH-IR marking noradrenergic neurons. (Scale bars=1 mm for light-sheet images, 50 µm for confocal images)

### C1QL3-2HA expression in the retina

While our prior work identified *C1ql3* expression in a subset of non-melanopsin-expressing retinal ganglion cells (RGCs) (Chew et al., 2017), our current analysis now reveals C1QL3-2HA expression in additional retinal layers and cell types, suggesting a broader role in visual signal processing. In the outer retina, we observed robust HA-IR labeling in the inner segment (IS) region of photoreceptors as well as in the outer plexiform layer (OPL), where photoreceptor synaptic terminals reside (Fig. 11A). Within the OPL, the HA signal appeared punctate and synaptically localized, consistent with a role for C1QL3 in photoreceptor output. This layer showed the most intense HA immunoreactivity in the retina, underscoring its potential importance in the early stages of light signal transduction.

**Figure 11:**
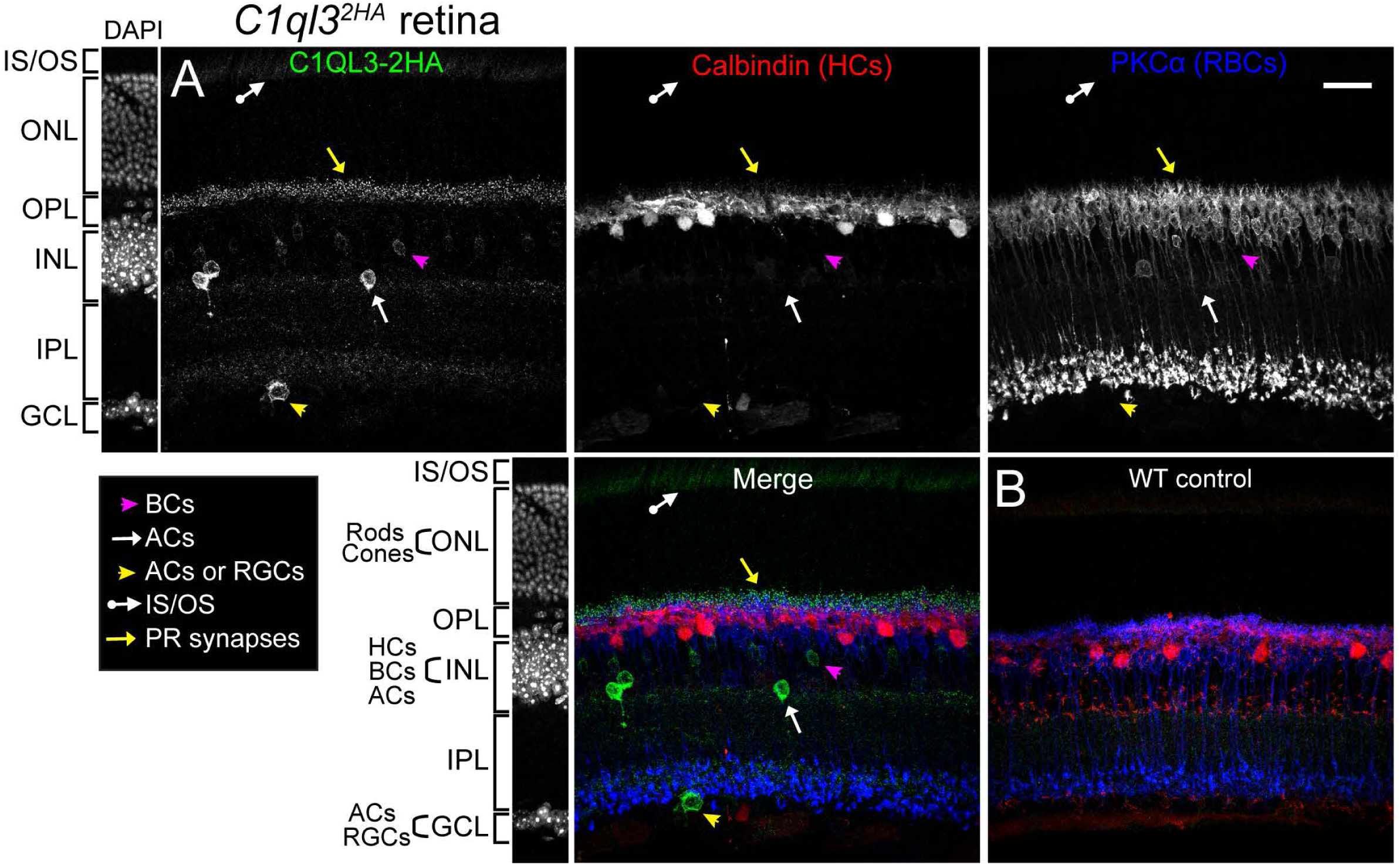
C**1**QL3**-2HA expression in the retina. (A)** Immunofluorescence for the indicated markers in a representative retina from an adult C1QL3^2HA^ mouse. Three distinct mice showed indistinguishable results. **(B)** WT mouse as a negative control for HA signal. Abbreviations: amacrine cells (ACs); bipolar cells (BCs); ganglion cell layer (GCL); horizontal cells (HCs); inner nuclear layer (INL); inner plexiform layer (IPL); photoreceptor inner segments/outer segments (IS/OS); outer nuclear layer (ONL); outer plexiform layer (OPL); photoreceptors (PRs); rod bipolar cells (RBCs); retinal ganglion cells (RGCs).

Moving inward, the inner nuclear layer (INL) displayed HA-labeled somata, consistent with those of cone bipolar cells (BCs). These were distinguished from rod BCs by the lack of colocalization with protein kinase C alpha (PKCα), a rod BC marker. Additional HA+ somas in the INL had the characteristic location and morphology of amacrine cells, further expanding the cellular reach of C1QL3. Importantly, double labeling with calbindin antibodies revealed no overlap between HA+ cells and horizontal cells, ruling out the expression of this protein in the inhibitory interneuron population.

In the ganglion cell layer (GCL), HA-IR cells likely included both displaced amacrine cells and RGCs. While the inner plexiform layer (IPL) contained some HA+ puncta, labeling here was much sparser compared to the intense signal in the OPL, suggesting that C1QL3 may play a more prominent role in the outer retinal synaptic circuitry. The complete absence of HA labeling in wild type retina (Fig. 11B) confirms the specificity of the signal. These findings align with recent single-cell RNA sequencing studies, which report strong *C1ql3* expression in photoreceptors, subsets of cone bipolar cells, amacrine cells, and RGCs, but low or absent expression in rod bipolar and horizontal cells (Shekhar et al., 2016; Tran et al., 2019; Yan et al., 2020). These results position C1QL3 as a molecular player in both the initial encoding of light stimuli by photoreceptors and the subsequent relay and refinement of visual signals by defined inner retinal circuits.

### C1QL3-2HA localization at mossy fiber synapses in the hippocampus

As noted previously, the density of C1QL3-2HA+ granule neurons in the dentate gyrus was so high that automated cell-counting algorithms failed to reliably quantify them. Consistent with light-sheet imaging, dense and robust HA expression was clearly evident in the granule cell layer (Fig. 12A–B). In addition, we detected a distinct population of HA+ neurons in the CA1 subregion (Fig. 12C) and a dimmer collection of HA+ cells and axons in the narrow CA2 zone (Fig. 12D).

**Figure 12:**
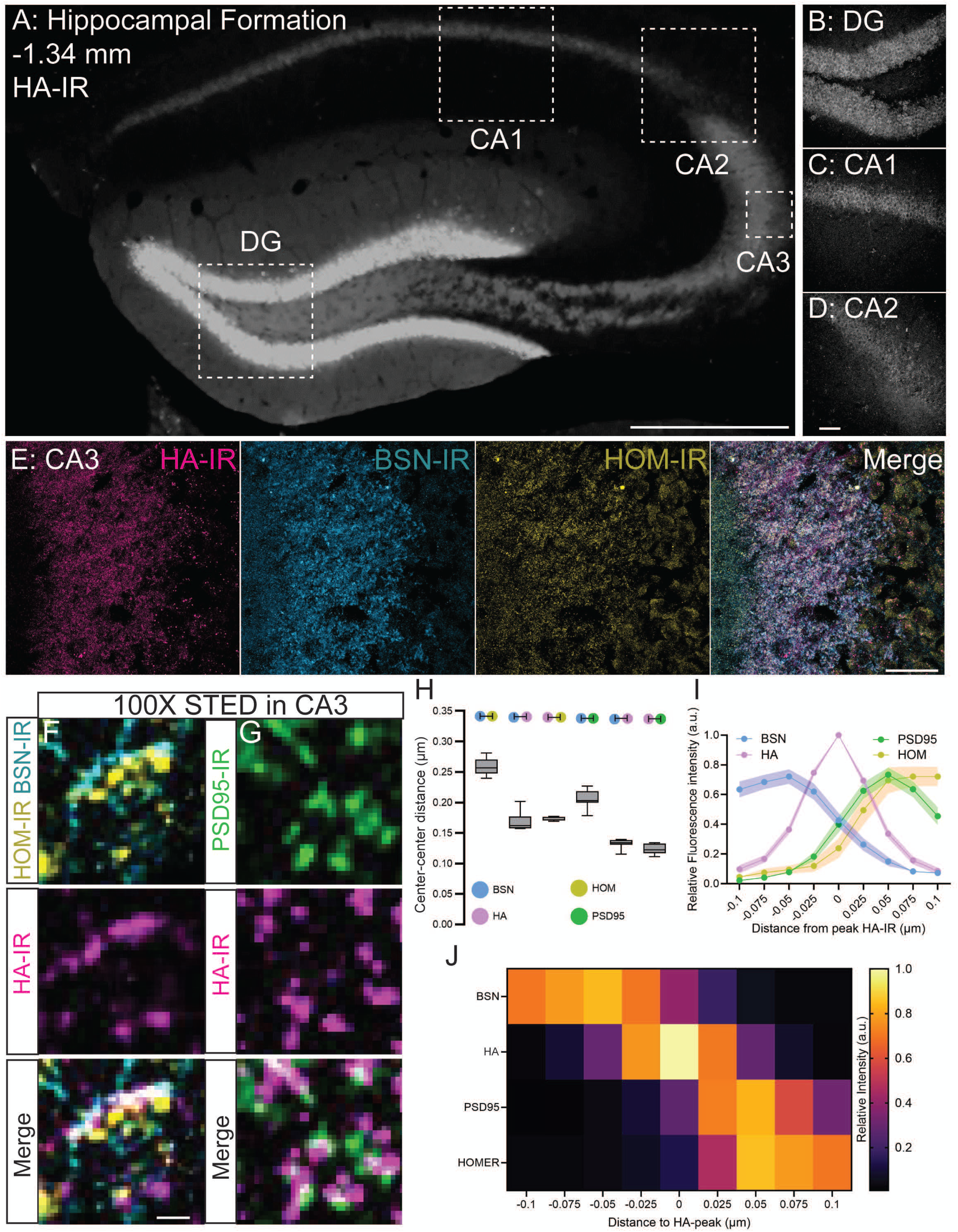
Subcellular localization of C1QL3-2HA at hippocampal mossy-fiber synapses. **A)** Fluorescent image (10X) of HA-IR in the hippocampus (scale bar=1 mm). **B)** Confocal images (40X) of HA-IR in the dentate gyrus (DG), **C)** CA1, and **D)** CA2. (scale bar=50 µm) **E)** Confocal images of axonal expression of HA-IR co-localized with presynaptic bassoon (BSN)-IR and postsynaptic homer (HOM)-IR in CA3 mossy fibers (100X, scale bar=50 µm). **F)** Representative 100X STED microscopy images of CA3 showing BSN-HOM-IR-defined synapses relative to C1QL3-2HA-IR. **G)** Representative 100X STED microscopy images of CA3 showing synapses defined by C1QL3-2HA-IR directly apposed to PSD95-IR. (scale bar=100 nm) **H)** Average center-to-center distances of all BSN, HA, and HOM-IR puncta under STED microscopy across six 100X ROIs within a single plane. Only HA-IR+ puncta within the average BSN-HOM distance were analyzed. **I)** Line graph of relative intensity through line profiles of distinct synapses defined by expression of BSN-IR, HA-IR, and HOM-IR as shown in panel F or HA-IR and PSD95-IR as shown in panel G, normalized to peak intensity for each, with zero set to the peak intensity of HA-IR at each synapse (n=41 for BSN, n=15 for HOM, n=38 for PSD95, n=53 for HA) **J)** Heatmap of relative fluorescence intensity of line profiles through distinct synapses defined by expression of BSN-IR, HA-IR, and HOM-IR as shown in panel F or HA-IR and PSD95-IR as shown in panel G, normalized to peak intensity for each, with zero set to the peak intensity of HA-IR at each synapse (n=41 for BSN, n=15 for HOM, n=38 for PSD95, n=53 for HA).

Most notably, the mossy fiber axons arising from granule cells and terminating in CA3 showed intense C1QL3-2HA labeling (Fig. 12E), consistent with earlier reports (Martinelli et al., 2016; Matsuda et al., 2016). This strong axonal signal in CA3 provided an ideal opportunity to leverage our KI line for high-resolution subcellular localization studies of C1QL3-2HA at synapses. We performed triple immunostaining for C1QL3-2HA, the presynaptic active zone marker bassoon (BSN), and the postsynaptic density proteins homer (HOM) and postsynaptic density protein 95 (PSD-95). In conventional confocal images, we observed robust colocalization of HA with both BSN and HOM in the CA3 region (Fig. 12E), as well as with BSN and PSD-95 (Supplementary Fig. S2A). To resolve the nanoscale architecture of these interactions, we employed stimulated emission depletion (STED) super-resolution microscopy. HA-IR consistently appeared to localize between paired BSN and HOM puncta, forming discrete, putative trans-synaptic alignments (Fig. 12F). Additional imaging with PSD-95 further confirmed that HA-C1QL3 puncta reside in close apposition to both presynaptic and postsynaptic compartments (Fig. 12G), suggesting a highly ordered spatial relationship across the synaptic cleft.

Quantitative STED analysis revealed that the center-to-center distances between BSN and HOM puncta matched previously published estimates from the mouse cortex (Akter et al., 2025). When we quantified the nearest-neighbor distances from C1QL3-2HA to either BSN and HOM or BSN and PSD95, we found the values to be nearly identical between the respective pre- and post-synaptic proteins in each experiment (Fig. 12H), strongly supporting a model in which C1QL3 bridges pre- and postsynaptic domains at the synaptic interface.

To further explore this synaptic topology, we performed fluorescence intensity line scans through individual HA+ puncta at putative mossy fiber synapses and aligned profiles across multiple sites (Supplementary Fig. S1, S2B). The resulting averaged traces revealed clear spatial segregation between presynaptic BSN, postsynaptic HOM and PSD-95, and C1QL3-2HA, which was consistently centered between them (Fig. 12I–J). These data support the hypothesis that C1QL3 functions as a trans-synaptic organizer, precisely positioned between active zones and postsynaptic densities (Matsuda et al., 2016), reminiscent of other synaptic bridging proteins (Dani et al., 2010; Trotter et al., 2019).

## DISCUSSION

In this study, we created and validated a new epitope-tagged C1QL3^2HA^ mouse line to map the native expression and subcellular localization of C1QL3 protein throughout the mouse brain and retina. Unlike previous methods that depended on mRNA profiling or fluorophore-based reporters (e.g. Martinelli et al., 2016) our model enables direct detection of endogenous C1QL3 protein without affecting its expression or function. Using this tool, we demonstrated that native C1QL3 assembles into high-molecular-weight oligomers of hexamers and 12-mers, consistent with our previous prediction using recombinant C1QL3 produced heterologous cells (Sticco et al., 2021). Whole-brain and retinal mapping revealed widespread and cell type-specific expression of C1QL3, including many previously unappreciated subcortical regions such as the midbrain raphe, VTA, regions of the hypothalamus (Figs. 5-6), as well as distinct cell populations identified by dual immunohistochemistry (Figs. 8–10). Furthermore, super-resolution STED imaging localized C1QL3 protein to the synaptic cleft of mossy fiber synapses in the hippocampus, consistent with previous work (Matsuda et al., 2016). C1QL3-2HA was situated between presynaptic and postsynaptic markers (Fig. 12), supporting a proposed role in trans-synaptic adhesion (Martinelli et al., 2016; Matsuda et al., 2016; Sticco et al., 2021). The C1QL3^2HA^ mouse thus provides a versatile and physiologically relevant model for investigating the molecular architecture and functional roles of C1QL3 in both the central nervous system and peripheral tissues where it is also expressed, such as the pancreas and adipose tissue (Chen et al., 2023; Gupta et al., 2018; Peña Palomino et al., 2023).

### Comparison with prior expression studies

Our C1QL3^2HA^ mouse line is a superior reporter compared to the previously developed *C1ql3*-mVenus mouse (Martinelli et al., 2016). Earlier analyses using *in situ* hybridization (ISH) from two groups (Iijima et al., 2010; Sigoillot et al., 2015) are consistent with our current findings and underscore the limitations of the mVenus reporter, which failed to detect multiple C1QL3-expressing populations. For instance, both ISH studies and our C1QL3^2HA^ mouse showed strong expression in the deep cerebellar nuclei, a region where mVenus expression was notably absent (Fig. 3Eii). We confirmed through qRT-PCR that whole-brain *C1ql3* mRNA levels are significantly lower in the *C1ql3*-mVenus mouse compared to our C1QL3^2HA^ mouse line and age-matched wild type controls, indicating hypomorphic expression in the reporter line (Fig. 2E).

However, this reduction likely varies by cell type: in the brainstem, for example, we observed many HA-IR+ cells that were not mVenus+, a difference that was less evident in forebrain areas (Fig. 3). We conclude that the C1QL3^2HA^ mouse provides a more accurate and consistent readout of endogenous C1QL3 protein expression.

A major advantage of our current study lies in the comprehensive, whole-brain analysis enabled by light-sheet microscopy, which offers superior spatial resolution and throughput compared to traditional histological methods. This approach uncovered many previously unreported populations of C1QL3-expressing neurons throughout the forebrain, midbrain, and hindbrain. Furthermore, the high-throughput capabilities of light-sheet imaging allowed us to estimate and compare the density of C1QL3-2HA+ cells across brain regions, revealing key anatomical patterns of expression (summarized in Figs. 5 and 6).

### Regional and laminar specificity across the CNS

Overall, the highest density of HA+ cells was observed in the cerebral cortex (Fig. 6A), but the expression pattern was far from uniform. In the neocortex, we found striking laminar specificity, with the most intense expression in layers 2/3 and 6, sparse and scattered labeling in layers 4 and 5, and complete absence from layer 1 (Fig. 7A–F). Counterstaining with calbindin revealed co-localization consistent with C1QL3 expression in some excitatory pyramidal neurons (Fig. 7H–I), in agreement with previous studies that described calbindin expression in a population of cortical projection neurons (DeFelipe, 1997). The observed laminar restriction of C1QL3-2HA indicates tightly regulated expression mechanisms that align with known functional compartmentalization in the neocortex. Layer 2/3 pyramidal neurons play a key role in intracortical processing and horizontal integration of sensory information (e.g. intratelencephalic projections), while layer 6 neurons are involved in feedback circuits to the thalamus and in modulating cortical excitability (Kim et al., 2014; Quiquempoix et al., 2018; Shipp, 2007; Yamashita et al., 2018). The enrichment of C1QL3-2HA in these layers, but less so in layer 5, which projects to subcortical targets, suggests that C1QL3 may support synaptic connectivity patterns specific to intracortical and corticothalamic communication.

Interestingly, this laminar pattern varied across different cortical regions. Neocortical areas showed the typical layered structure, while allocortical regions such as the hippocampus, entorhinal cortex, and insular and piriform cortices displayed more diffuse or regionally specific expression. Notably, C1QL3-2HA was broadly expressed in the dysgranular zones that mark the transitions between these cortical types (Fig. 5B–I). Our data suggest that C1QL3 might help organize the molecular structure of these integrative hubs. Similarly, in the superior colliculus (SC), C1QL3-2HA+ cells were limited to the superficial sensory layers, with no expression in the deeper motor layers or in the nearby inferior colliculus (Fig. 5H–I).

In the thalamus, we observed a notably discrete pattern of C1QL3-2HA expression that closely parallels the distribution of calretinin+ thalamic neurons (Fig 5. D-E; Viena et al., 2021), a relationship we confirmed through co-labeling experiments (Fig. 8B–C). Several HA+ thalamic nuclei, including the paraventricular (PVT), central medial (CM), and interanterodorsal (IAD) nuclei, are part of the intralaminar and midline thalamic group known to project diffusely to widespread cortical areas. These thalamic regions have been linked to arousal, salience processing, sleep-wake regulation, and the coordination of cortical states (Clascá et al., 2012; Harris et al., 2019; Hunnicutt et al., 2014). The presence of C1QL3 in these neurons suggests it may play a role in maintaining the structural integrity or fine-tuning the synapses of long-range thalamocortical projection systems that support global cortical functions.

Given C1QL3’s role as a secreted synaptic organizer, its selective expression in these projection neurons suggests it may facilitate specific synaptic connectivity or long-term maintenance or synaptic strength of thalamocortical circuits that are functionally distinct from the primary sensory relay thalamus. Additionally, the non-overlapping expression patterns of C1QL3-2HA and parvalbumin as noted in the internal globus pallidus but not in the striatum or thalamus (Fig.8D) indicate cell type-specific functions that could underlie C1QL3’s roles in connectivity and plasticity within distributed brain networks.

Together, these data demonstrate how the C1QL3^2HA^ mouse enables us to distinguish regionally and laminar-specific populations that were previously undetectable or unclear. They suggest that C1QL3 may be expressed in circuits with distinct synaptic functions, whether for local intracortical processing, long-distance corticothalamic feedback, or broad modulatory projections, and may help define or maintain synaptic partnerships within these specific anatomical areas.

### C1QL3 in inhibitory and monoaminergic populations

While most C1QL3-expressing neurons are excitatory, our data confirm and expand upon earlier findings indicating that C1QL3 is also present in specific inhibitory cell types.

Previously, we identified arginine vasopressin (AVP) and vasoactive intestinal peptide (VIP) neurons in the suprachiasmatic nucleus (SCN) as inhibitory populations that express C1QL3 (Chew et al., 2017). Here, we additionally identified Golgi interneurons in the cerebellar granule cell layer as a previously unreported inhibitory cell type that expresses C1QL3 (Fig. 9H). These interneurons are critical regulators of cerebellar glomerular function, modulating granule cell excitability and contributing to motor coordination and sensorimotor integration (Consalez et al., 2021; Lee et al., 2023; Okada et al., 2022). We also found C1QL3 expression in parvalbumin neurons of the GPi, another inhibitory population (Wallace et al., 2017), although HA-IR cell bodies were additionally present in non PV+ cell types, suggesting C1QL3 expression by additional cell types in the region that may warrant future study.

Whether C1QL3 functions similarly in inhibitory neurons as it does in excitatory projection cells is unknown. It is possible that in inhibitory populations, C1QL3 plays distinct roles, including non-synaptic functions. Supporting this idea, we recently showed that the closely related family member C1QL1 regulates the differentiation of oligodendrocyte precursor cells (Altunay et al., 2025). Furthermore, outside the CNS, C1QL3 is involved in various physiological processes, including insulin secretion from pancreatic β-cells and the regulation of systemic glucose and lipid metabolism (Chen et al., 2023; Gupta et al., 2018; Peña Palomino et al., 2023). These findings underscore the possibility that C1QL3’s function may be cell type- and tissue-specific, and not strictly limited to synaptic adhesion, and furthermore how the C1QL3^2HA^ mouse will be a key tool in dissecting the likely varied role of C1QL3 across different cell types.

Besides inhibitory neurons, we observed C1QL3 expression in several neuromodulatory groups that fall outside of usual excitatory/inhibitory categories. C1QL3-2HA was highly expressed in select serotonergic neurons of the raphe magnus and dorsal raphe (Fig. 9B–C), as well as in a sparse subgroup of dopaminergic neurons in the ventral tegmental area (VTA) (Fig. 10B), but notably not in noradrenergic neurons of the locus coeruleus. These cell groups are well-established regulators of behavioral state, arousal, and emotion. The specific expression of C1QL3 in only a subset of VTA dopaminergic neurons is particularly noteworthy and supports increasing evidence that this region contains molecularly varied neuronal subtypes. Many VTA neurons are now known to co-release GABA or glutamate (Kawano et al., 2006; Margolis et al., 2012; Miranda-Barrientos et al., 2021; Morales & Margolis, 2017; Tritsch et al., 2012), raising the possibility that C1QL3 contributes to synaptic specificity or output modulation in these functionally distinct subcircuits.

### Retinal expression and cell type-specificity

In the retina, we observed C1QL3-2HA expression in photoreceptors, cone bipolar cells, amacrine cells, and RGCs, consistent with prior transcriptomic studies (Shekhar et al., 2016; Tran et al., 2019; Yan et al., 2020), but contrasting the underreported subset of cells that express *C1ql3* in the retina of the mVenus reporter mouse (Chew et al., 2017). HA labeling was especially intense in the photoreceptor inner segments (IS) and outer plexiform layer (OPL), where it appeared punctate and likely localized to synaptic terminals (Fig. 11A). Within the inner nuclear layer (INL), HA+ somata were observed in bipolar cells, which we determined to be cone bipolar cells based on a lack of colocalization with PKCα, a rod bipolar marker. Additional HA+ somata with amacrine-like morphology were also present in the INL, and in the ganglion cell layer (GCL), where they likely represent displaced amacrine cells and RGCs. In contrast, HA labeling was absent in horizontal cells, as confirmed by double-staining with calbindin (Fig11A). Interestingly, labeling in the inner plexiform layer (IPL) was sparse compared to the OPL, suggesting a more prominent presynaptic role in photoreceptors than in inner retinal circuitry. The specificity of the signal was validated by the absence of HA labeling in WT retina (Fig. 11B). Notably, C1QL3-2HA labeling partially overlaps with, but is spatially and cellularly distinct from that of C1QL1-2HA, which is enriched in horizontal cells and the IPL (Cheung et al., 2024), suggesting complementary functions. Given the known interaction between C1QL3 and the adhesion GPCR ADGRB3, expressed in multiple bipolar cell types (Shekhar et al., 2016), it is plausible that C1QL3 is secreted from photoreceptors and acts trans-synaptically to modulate bipolar cell synapses, similar to its role in central brain circuits (Martinelli et al., 2016; Matsuda et al., 2016; Wang et al., 2020).

### Synaptic localization and trans-synaptic positioning

Using super-resolution STED microscopy, we precisely localized C1QL3 at mossy fiber synapses in the hippocampal CA3 region (Fig.12). Previous work using a non-commercially available C1QL3 antibody qualitatively showed at super-resolution using SIM microscopy that C1QL3 was localized between BSN and PSD95 puncta in CA3 (Matsuda et al., 2016). We expanded on this work using our C1QL3^2HA^ mouse to localize HA-IR between presynaptic bassoon (BSN) and postsynaptic homer (HOM) and PSD95 using STED microscopy, which has improved resolution from SIM (Wegel et al., 2016). In this study, C1QL3-2HA consistently appeared equidistant between presynaptic BSN and postsynaptic HOM and PSD-95 (Fig.12F, G). To quantify this relationship, we measured the center-to-center distances between BSN and either HOM or PSD95 puncta in single-plane 100x STED images, obtaining values consistent with prior high-resolution estimates of synaptic cleft width (Fig. 12H, Akter et al., 2025). When we calculated the nearest-neighbor distances between C1QL3-2HA puncta and either BSN and HOM or BSN and HOM, the values were nearly identical between HA and the respective pre-and post-synaptic protein in each experiment, reinforcing the notion that C1QL3 localizes relatively symmetrically within the cleft at mossy fiber synapses (Dani et al., 2010). Line scan analysis of fluorescence intensity through individual HA+ synapses further confirmed this spatial arrangement: averaged intensity plots showed clear separation between pre- and post-synaptic markers with C1QL3-2HA centered between them (Fig. 12I–J). This supports a model in which C1QL3 functions as a trans-synaptic organizer, spanning the synaptic cleft (Martinelli et al., 2016; Matsuda et al., 2016; Sticco et al., 2021), similar to what has been proposed for cerebellins, neurexins, and other adhesion proteins (Südhof, 2023; Trotter et al., 2019). These findings provide the most precise localization to date of endogenous C1QL3 at hippocampal synapses and highlight the utility of the C1QL3^2HA^ mice for dissecting synaptic nano-architecture *in vivo*. This model offers a unique platform for future biochemical, imaging, and structural studies aimed at defining the molecular composition and dynamic behavior of C1QL3-containing synaptic adhesion complexes.

### Biochemical opportunities and structural considerations

The addition of two HA tags to endogenously expressed C1QL3 enables direct extraction and purification of the protein from native tissue, allowing for robust biochemical analyses, including immunoprecipitation and co-immunoprecipitation of C1QL3-containing complexes. We and others have previously demonstrated that C1QL3 can form higher-order oligomers *in vitro* (Bolliger et al., 2011; Iijima et al., 2010; Sticco et al., 2021; Wei et al., 2011). In this study, we confirmed that native C1QL3 assembles into hexamers and dodecamers *in vivo*. This C1QL3^2HA^ mouse model allows us, for the first time, to examine the biochemical, structural, and functional properties of endogenous C1QL3 and its tissue-specific assembly states. C1QL3 oligomeric states may affect binding avidity, receptor clustering, and trans-synaptic specificity, which are crucial for establishing or maintaining different types of synapses. Combining this model with structural techniques such as crosslinking mass spectrometry or electron microscopy could enable the reconstruction of native synaptic C1QL3 assemblies *in situ* and provide new insights into the effector mechanisms of synaptic plasticity.

Importantly, our STED imaging data underscore the value of our novel mouse line for future studies on C1QL3’s binding partners, including neurexins and kainate receptors (Matsuda et al., 2016), the adhesion GPCR ADGRB3 (Bolliger et al., 2011), and the neuronal pentraxins (Sticco et al., 2021). The C1QL3^2HA^ mouse provides an ideal platform for dissecting these interactions biochemically and anatomically. Given the overlapping and complementary expression patterns of C1QL3 with other C1q superfamily members such as C1QL1, C1QL2, and cerebellins, future studies may also explore competitive or cooperative binding mechanisms that converge on shared postsynaptic targets and modulate synapse identity, plasticity, or stability.

### Behavioral and circuit-level applications

In our previous studies of C1QL3 function, we identified a range of behavioral deficits in constitutive *C1ql3* knockout (*C1ql3^-/-^*) mice, including impairments in cognitive flexibility, motor learning, and emotionally salient memories (Caro et al., 2025; Chew et al., 2017; Martinelli et al., 2016). The brain-wide expression patterns revealed in the current study now provide a framework to link these phenotypes to specific neural populations and circuits.

For example, we identified strong C1QL3-2HA expression in cerebellar Golgi interneurons and neurons of the deep cerebellar nuclei (Fig. 3E–I, 9E), both of which are central to cerebellar circuits regulating motor learning (Consalez et al., 2021; Okada et al., 2022).

These populations are promising candidates for mediating the motor deficits observed in *C1ql3^–/–^* mice. Similarly, we identified C1QL3-2HA+ serotonergic neurons in the raphe nuclei (Fig. 9B–C), as well as a sparse subpopulation of dopaminergic neurons in the VTA (Fig. 10B), suggesting that C1QL3 may contribute to motivated behaviors through specific neuromodulatory subcircuits. Other regions with strong C1QL3-2HA expression form parts of well-established behavioral pathways. For example, several midline thalamic nuclei that relay information between the medial prefrontal cortex and the amygdala, regions central to fear processing, also express C1QL3 (Penzo & Gao, 2021). This supports earlier findings that *C1ql3* deletion in the amygdala impairs fear memory (Martinelli et al., 2016). Additionally, we confirm high expression in the (AON), where C1QL3 was shown to be a key player in olfactory memory (Wang et al., 2020). Finally, in the claustrum, a region we found to exhibit widespread C1QL3-2HA expression, is known to coordinate cortical-subcortical integration for sensory and attentional processing (Nikolenko et al., 2021; Remedios et al., 2010). Taken together, these observations suggest that C1QL3 may mediate complex behaviors through its role in multiple nodes of distributed functional networks. Future studies will be needed to determine whether loss of *C1ql3* from specific regions or circuits is sufficient to produce the behavioral phenotypes observed in global knockout mice.

### Limitations

This study has several technical limitations. First, detecting C1QL3-2HA requires immunohistochemistry, as the HA epitope is not intrinsically fluorescent. Antigen retrieval was necessary to visualize synaptic labeling, which can affect tissue morphology and limit multiplexing with some antibodies. Second, densely packed cell regions, such as the dentate gyrus, presented challenges for automated cell quantification, occasionally underestimating actual numbers. Minor misalignments in our light-sheet data during anatomical registration to the Allen Brain Atlas may also impact subregion-level quantifications, particularly for small or adjacent subnuclei. Despite these limitations, the detailed expression mapping and validation provided here offer a solid basis for future targeted studies.

### Conclusions

Overall, our results establish the C1QL3^2HA^ mouse model as a valuable tool for dissecting the molecular and anatomical basis of C1QL3 expression, structure, and function. We demonstrate that our model allows for high-resolution mapping, precise targeting, and biochemical access to C1QL3-expressing neurons across various brain systems involved in cognition, vision, perception, arousal, and autonomic regulation. Its specificity makes it particularly useful for future studies on synaptic organization, connectivity, and dysfunction in neurological and psychiatric disorders linked to disrupted circuit architecture, such as epilepsy, schizophrenia, and autism spectrum disorders.

## Data availability

The data that support the findings of this study are available from the corresponding author upon reasonable request.

## Acknowledgements

We gratefully acknowledge all members of the Jackson, Martinelli, Ressl, and Lee labs for support, assistance and helpful discussions, K. Lowther and D. Kabac at The Center for Mouse Genome Modification (CMGM) at UConn Health for creation of the *C1ql3^2HA^* knock-in allele and O. Knuth for genotyping support and mouse colony management. Many thanks to A. Shankardass of LifeCanvas Technology for managing our tissue-clearing and light-sheet-imaging workflow. Thanks to M. Hruska for advice on STED imaging, E. Wainman for assistance with the Allen Brain MERFISH database and light-sheet quantification and to P. Peña Palomino for assistance with the design of the knock-in construct. Project was supported by the National Institutes of Health (NIH) National Research Service Award (NRSA) fellowship F31HL165896 (NHLBI to WPA) and NIH grants R01MH112739 (NIMH to ACJ), R01NS131664 (NINDS to ACJ and DCM), R03DC019290 (to DCM), R01EY026817, R03TR005086 (to AL), NIH Shared Instrumentation grants S10OD016435 (to A. Nishiyama) S10OD023618 (to CO) and UT Austin start-up funds 57555R6T and R03TR004494 (to SR). We also thank the Connecticut Institute for the Brain and Cognitive Sciences (IBACS) for a graduate summer fellowship (to WPA).

---

**Supplementary Figure S1.**
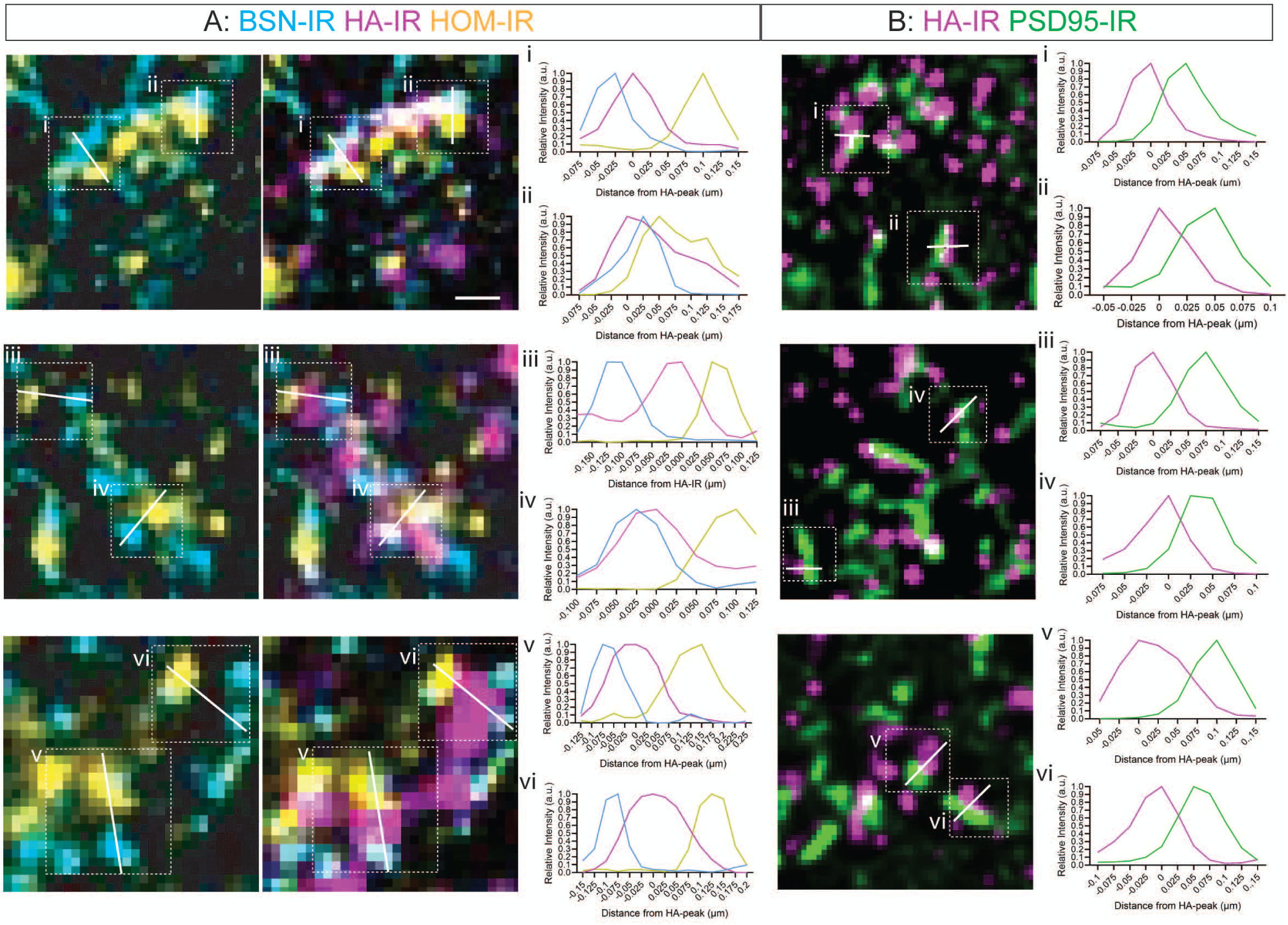
: Example putative mossy fiber synapses from STED experiments. **A)** Individual examples of line scans in STED images (100X) through putative mossy fiber synapses as defined by BSN-HOM appositions which also contained HA-IR in CA3 mossy fibers (left) with relative fluorescence intensity of each quantified (right). **B)** Examples of line scans in STED images (100X) through putative synapses as defined by HA-PSD95 appositions in in CA3 mossy fibers (left) with relative fluorescence intensity of each quantified (right). (All scale bars=100 nm)

**Supplementary Figure S2.**
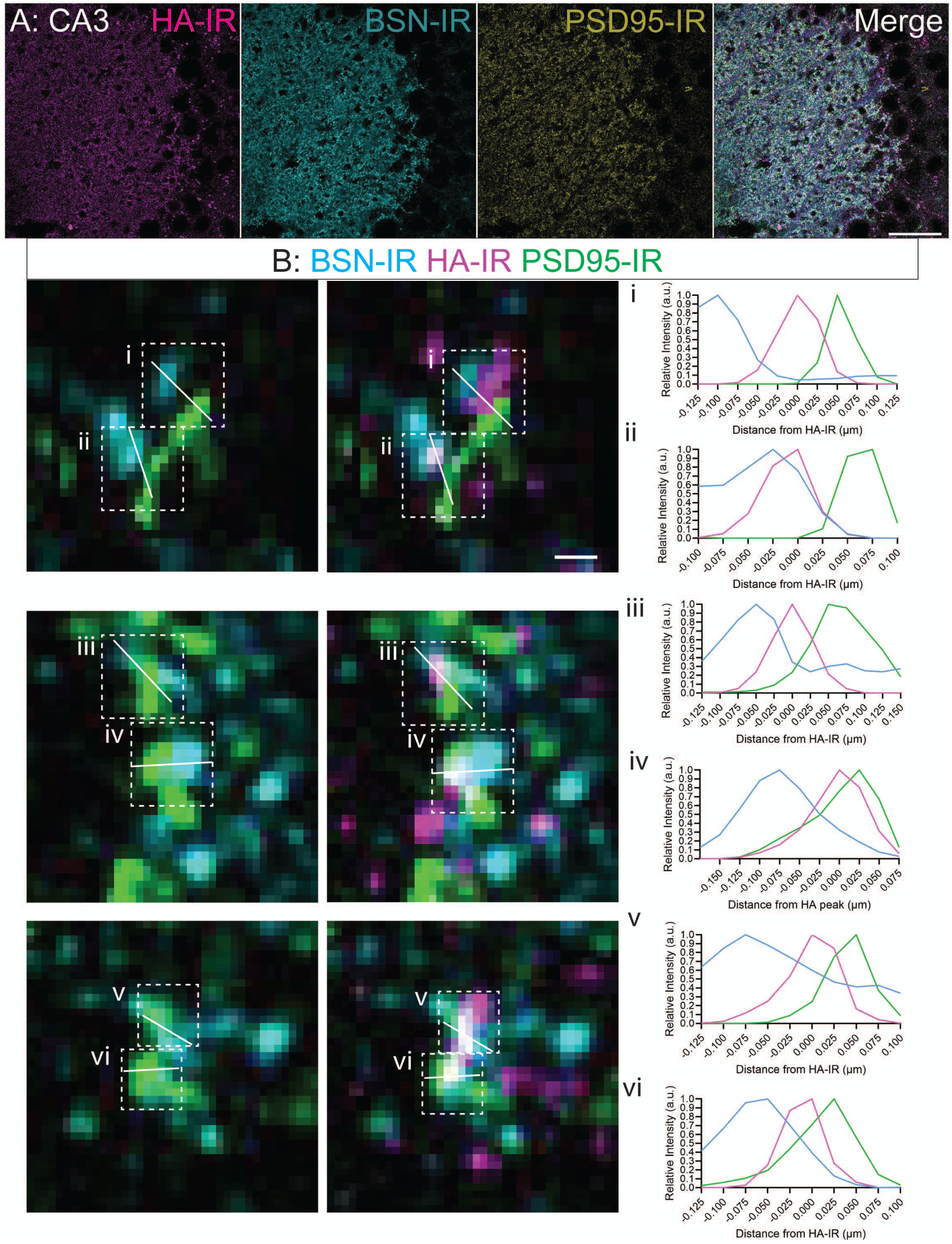
: BSN-HA-PSD95 defined mossy fiber synapses in CA3 of C1QL3^2HA^ mice. **A)** Confocal images (100X) of axonal expression of HA-IR co-localized with presynaptic BSN-IR and post-synaptic PSD95-IR in CA3 mossy fibers. (scale bar=50 µm) **B)** Examples of line scans in STED images (100X) through putative synapses as defined by BSN-PSD95 appositions which also contained HA-IR in CA3 mossy fibers (left) with relative fluorescence intensity of each quantified (right). (scale bar=100 nm)

